# A soft active matter models explains spiral epithelial cell migration on in-vivo corneas

**DOI:** 10.1101/2024.08.14.608002

**Authors:** Kaja Kostanjevec, Rastko Sknepnek, Jon Martin Collinson, Silke Henkes

## Abstract

The mammalian cornea constantly regenerates its outer epithelial layer. Cells lost by abrasion are replaced by division of both corneal epithelial cells and stem cell populations around the corneal periphery, the limbus. Limbal-derived epithelial cells migrate into the cornea, maintaining equal rates of cell loss and replacement (the ‘XYZ hypothesis’). This process produces a striking stable spiral cell motion pattern across the corneal surface, with a central vortex. Here, we show that this spiral pattern can be explained by the interplay of limbus position, cell division, extrusion, and collective cell migration along the curved corneal surface. Using dissected LacZ mosaic murine corneas, we inferred the surface flow field by following stripe edges, revealing a tightening spiral. To explain these flow fields, we developed a cell-level in silico model treating corneal epithelial cells as soft, self-propelled particles with density-dependent proliferation and extrusion rates, and noisy alignment of migration direction. Even without global guidance cues, the model predicted stripes and spirals closely recapitulating experiment. A complementary continuum description generalised the XYZ hypothesis. Spiral formation was robust to curvature changes, but not topology, and sensitive to limbal stem cells and flocking alignment, showing how swarm physics on curved surfaces can explain tissue-scale biological processes.

## Introduction

The transparent cornea, the most anterior structure of the vertebrate eye, consists of a hypocellular, collagenous stroma covered by a stratified epithelium, resting on an inner endothelial monolayer (***Forrester et al., 2020***). The corneal epithelium, which is contiguous with the conjunctival epithelium that covers the rest of the anterior of the eye and the inner edge of the eyelids, is a paradigm model of tissue maintenance by stem cells (***Cotsarelis et al., 1989***; ***Lavker et al., 1991***; ***Lehrer et al., 1998***). Cells are constantly lost from the corneal surface by abrasion and must be replaced. Multiple lines of evidence have shown that the stem cells that maintain the corneal epithelium reside in a narrow ring at the edge of the cornea, at the boundary between the corneal and conjunctival stem cells – the limbus (***Cotsarelis et al., 1989***; ***Richardson et al., 2017***). Limbal epithelial stem cells themselves are organised as an outer ring of slow-cycling, label-retaining cells that most likely contribute to corneal regeneration after wounding, and an inner layer of rapidly dividing cells that contribute to normal corneal homeostasis (***Altshuler et al., 2021***; ***Farrelly et al., 2021***). Stem cells in the basal layer of the limbal epithelium divide and produce proliferative, undifferentiated ‘transit-amplifying’ (TA) cells that migrate into the corneal epithelium to replace those lost during normal life. Studies of human ‘hurricane’ keratopathies and mosaic patterns of cell staining in genetically tagged rodents, frogs and fish have shown that the tracks of migrating cells show a remarkable, regular, radial pattern in the uninjured corneal epithelium, usually with a central vortex (***Bron and Tripathi, 1973***; ***Dua et al., 1993***; ***Collinson et al., 2002, 2004***; ***Nagasaki and Zhao, 2003***; ***Iannaccone et al., 2012***; ***Di Girolamo et al., 2015***; ***Nejad et al., 2015***). A similar pattern develops during growth of the retina of the eye (***Tsingos et al., 2019***). The pinwheel patterns of cell staining observed suggest that migration is highly regular and potentially globally coordinated. The vortices of cell motion patterns observed at the corneal epithelial centre resemble a logarithmic spiral and lend themselves to mathematical analysis (***Iannaccone et al., 2012***). Modelling suggests the spiralling may correlate with tension vectors due to intraocular pressure (***Nejad et al., 2015***), but there are no empirical studies suggesting how it might occur in vivo.

The drivers for directed cell migration across the ocular surface are poorly known. Various mechanisms have been suggested, including chemorepulsion from vascular components at the limbus, population pressure from the highly proliferative peripheral corneal epithelium, biased cell loss near the corneal centre, electrical currents caused by ion leakage through abrasion, and contact-mediated cues (***Bron and Tripathi, 1973***; ***Buck, 1985***; ***Sharma and Coles, 1989***; ***Lemp and Mathers, 1989***; ***Lavker et al., 1991***; ***Wolosin et al., 2000***; ***McCaig et al., 2005***; ***Foster et al., 2014***). Patterns of cell migration are known to be disrupted, at least temporarily, by wounding, and in disease states (***Collinson et al., 2004***; ***Mort et al., 2009***; ***Richardson et al., 2018***). The planar polarity protein, Vangl2, is required for normal in vivo patterns of cell migration, suggesting aligned cell-cell interactions (***Findlay et al., 2016***). Corneal epithelial cell migration can be directed in a contact-mediated manner by substrate topography, and an integrin-mediated, cAMP-dependent interaction with their basement membrane may maintain some degree of radial orientation of migrating cells in vitro, but is not sufficient to push all cells in a centripetal direction (***Walczysko et al., 2016***). A major centripetal driver may be a step-change increase in substrate compliance at the limbal-corneal boundary (from soft to hard substrate), causing limbal cells to make an initial durotactic movement into the cornea (***Walczysko et al., 2016***; ***Gouveia et al., 2019***).

However, it has not been shown experimentally whether these genetic couplings or mechanical and extrinsic directional cues would be enough to produce stable patterns of long-distance migration in vivo. Understanding the coordinated, highly patterned, constrained collective migration patterns of the corneal epithelial cell sheet in vivo is a complex three-dimensional biological problem requiring an integrated, systems-based resolution that goes beyond the individual contributing elements.

An in silico model of in vivo ocular surface maintenance that produces radial striping is not conceptually difficult, and coordinate-based models based on cell proliferation dynamics and/or cell density have been proposed (***Lobo et al., 2016***; ***Moraki et al., 2019***). They, however, have unsatisfactory aspects from a biological and predictive perspective. A good model needs to explain, not simply describe, both the formation and maintenance of stripes and the central spiral of cell movement, within a confluent cell sheet. It needs to be formulated in three dimensions to account for the curvature of the cornea, to be quantitative and scaleable, in both size and simulation times, and to apply to corneas of different shapes; importantly, it needs realistic local dynamics of physical cell-cell interactions, incorporate active cell migration and planar alignment, and to make no assumptions about cell behaviour or guidance that are not rooted in observations. A realistic in silico model of corneal wounding and maintenance using the cellular Potts model was recently proposed (***Vanin et al., 2025***), for a local, vertical slice through the tissue. To link the space and time scales from individual cells up to the full corneal motion, however, needs more conceptually:

The physics of active or living matter seeks to understand the collective properties of individually motile agents, such as birds, fish, bacteria or eukaryotic cells (***Vicsek and Zafeiris, 2012***; ***Marchetti et al., 2013***; ***Needleman and Dogic, 2017***). One of its hallmarks is a state of collective migration, or ‘flocking’ that generally appears when agents *locally* coordinate their motion, without the need for a leader or guidance cue (***Vicsek et al., 1995***; ***Giardina, 2008***). Such flocking collectives where the directions of motion are all aligned with each other, are known as polar active materials. The apparent flocking behaviour of cells across the corneal surface, together with known behaviours of individual cells, such as self-propulsion, local coordination, contact-mediated cell proliferation and differentiation, leads us to hypothesise that a collective migration model rooted in the physics of soft active matter (***Alert and Trepat, 2020***) can explain the observed in vivo migration patterns.

Within the framework of active matter, it has become clear that the system’s topology plays a prominent role. For example, a sphere is a closed surface with no holes, with an associated topological invariant, the Euler characteristic *χ* = 2, loosely speaking, a number that describes an object’s shape regardless of how it is bent. A mathematical consequence is that the velocity profile of any flow confined to the surface of a sphere has to vanish at least at one point. Flocks, therefore, cannot maintain fully aligned velocities everywhere on the surface, and instead create points with locally mismatched directions. Near such ‘singular points’, if one loops over a contour encompassing it, the flow field winds an integer number of times (***Do Carmo, 2016***). Such points are known as *topological defects* with *topological charge c* (***Mermin, 1979***). Pictorially, +1 defects are circular objects corresponding to inward, outward or spiral flows around the defect core, while −1 defects have four-leaf clover symmetry, with flow inwards along two directions and outwards along the orthogonal ones (***Alexander et al., 2012***). For a brief overview of topological charges, please see Box 1. The Poincaré-Hopf theorem requires that the total topological charge equals the Euler number, i.e. ∑_*i*_ *c*_*i*_ = *χ* (***Do Carmo, 2016***). Flocks of active particles on a sphere then typically generate patterns consisting of a circling band and two vortex-like +1 topological defects at the poles (***Sknepnek and Henkes, 2015***; ***Shankar et al., 2017***; ***Hsu et al., 2022***).

The cornea, with the limbal boundary, is a spherical cap, i.e. it has the same topology as a disk. In this topology, with Euler characteristic *χ* = 1, if the flows close to the boundary are uniform as for the cornea, we expect at least one topological defect with charge +1 (***Stein, 1979***). We propose that this is the vortex pattern corresponding to the central spiral of the cornea. Indeed, in at least two in vitro collective cell system with a disk topology, such a spiral defect was described and linked to topology (***Guillamat et al., 2022***; ***Lång et al., 2024***). Pictorially, the migration pattern for the cornea is akin to water flowing in from a source and then circling down a sink, except that we can create and extrude cells everywhere within the sheet. Additionally, biological development can also mould the surfaces themselves and then generate deformations that couple to topological defects – as has recently been shown for hydra (***Maroudas-Sacks et al., 2021***), and argued to play an important role for organoids (***Rozman et al., 2020***).

In this paper, we build a quantitative model of spiral flocking on the cornea. We previously showed that the swirling migration patterns of confluent corneal epithelial cell monolayers in vitro can be understood using a model of active matter at high densities (***Henkes et al., 2020***), and that one can include mechanical coordination between cells as polar alignment (***Saraswathibhatla et al., 2021***). The model treats each cell as a soft particle, able to self-propel, align its planar polarity with, and exert elastic forces on its neighbours. These features are accounted for by a set of equations of motion that evolve in time to simulate swirling spatiotemporally correlated patterns of cell motion as observed in in vitro experiments. The model is quantitative, and its parameters can be directly derived from experimental observation.

Here, we extend this model of corneal epithelial cell migration to a three-dimensional constrained geometry that recapitulates the mouse cornea, with stem cells at the limbus and proliferating TA cells within the cornea. The model produces radial striping flow patterns with a central spiral representing a +1 topological defect that replicates the spiral flow patterns of cell migration observed in vivo. We link flocking spiral formation to effective resurfacing of the cornea, and reformulate the XYZ hypothesis of corneal maintenance (***Thoft and Friend, 1983***) in physical terms. Additionally, using simulations of the model in different conditions, we are able to show that spiral formation is robust to changes in corneal shape, as long as the *χ* = 1 disk-like topology is preserved.

### Box 1.

**A brief overview of topological charges**

**Box 1 Figure 1.**
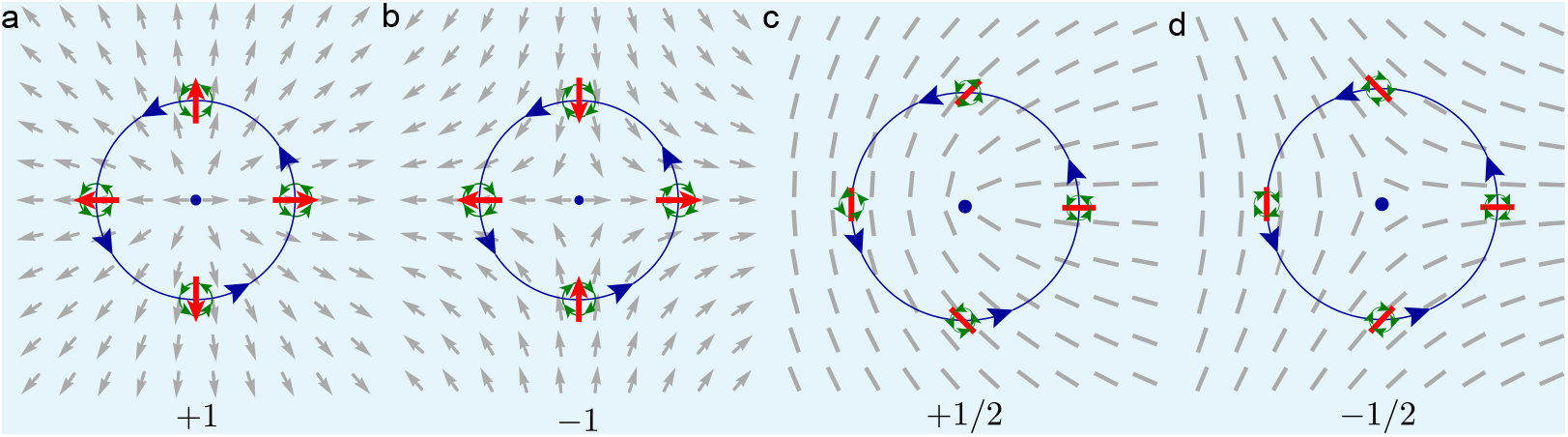
Topological defects in two-dimensional polar and nematic fields. Grey arrows denote polar vectors, while grey lines denote nematic directions with head–tail symmetry. **a** Looping counter-clockwise (blue circle) around the defect core (blue dot), the polar vectors rotate counterclockwise by 2*π*, giving a topological charge of +1. **b** Looping counterclockwise, the vectors rotate clockwise by 2*π*, corresponding to a charge of −1. **c** In a nematic field, head–tail symmetry allows a half-turn of the field; the vectors rotate by *π*, giving a charge of 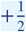. **d** The same configuration as in panel c but with clockwise rotation, yielding a charge of 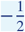. A uniform rotation of all vectors leaves the topological charge unchanged, making all spirals +1 topological defects.

A vector field, such a cell migration forces and velocities, assigns a direction to every point on a surface. In some systems, such as the nematic liquid crystals that model materials made out of long thin objects, the field has head–tail symmetry, so reversing a direction has no physical effect. Distortions of two-dimensional vector fields fall into two classes. *Tilts* are smooth variations that can be removed by a global rotation of all vectors. *Topological defects* are localised singularities with a core where the field direction is ill-defined and cannot be eliminated by any smooth reorientation.

The defining property of a defect is the winding of the field around its core. Traversing a closed loop counterclockwise, the surrounding vectors rotate by a total angle of 2*nπ* (or −2*nπ*), giving a topological charge of +*n* (or −*n*). For polar fields, *n* is an integer, while nematic fields admit half-integer charges due to head–tail symmetry.

On a closed surface without boundaries, the sum of all defect charges is fixed by the Euler characteristic *χ* = 2 − 2*g*, where *g* is the number of holes in the surface. For example, a sphere (*g* = 0) requires a total charge of 2, whereas a torus (*g* = 1) has zero net charge. For surfaces with boundaries, the total interior charge depends on the boundary conditions; for a disk where the field at the boundary is either perpendicular or tangent to it, known as perpendicular or tangent anchoring respectively, the net charge is +1.

In contrast, spiral formation does not happen at all without cell-cell alignment and is severely perturbed when the limbal stem cell niche is absent. We are thus able to demonstrate the central role of flocking, confinement and curved geometry in corneal patterning. This model represents the first comprehensive, predictive tool for understanding the maintenance of the full ocular surface in health and disease.

## Results

### Experimental flow fields

Figure 1a shows a representative X-linked LacZ-transgenic adult female mouse cornea, with a pin-wheel spiral pattern of β-galactosidase mosaicism due to the flow of cells from patches of blue and white stem cells at the periphery (limbus) to the corneal centre throughout adult life (***Collinson et al., 2002***). The patterns of cell migration are reflected by the projection of sensory axons between the cells in the basal layer, which appear to largely follow the flow of epithelial cells (Figure 1b), and form together with the corneal epithelium during development. Both axons and cells form a smooth flow pattern with a vortex at the corneal centre. This vortex is an example of a topological defect (i.e. a point where the orientational order of cell velocity vectors vanishes) with a +1 topological charge, as described in Box 1, and shown in Box 1 Figure 1a. We hypothesise that the flow pattern and the central topological defect can be explained as an emergent behaviour of a collective of self-propelled epithelial cells closely packed on a hemisphere. As a differential hypothesis, epithelial cells on a substrate have been described as an active nematic material (***Saw et al., 2017***; ***Balasubramaniam et al., 2021***), where elongated cells or the stresses of the cells pulling on each other generate topological defects with ±1/2 charges, as again described in Box 1. Despite the compatible elongated shape of the axons, we do not observe such defects, which for a +1/2 defect resemble a fold in a material made of long, thin objects (see Box 1 Figure 1c-d). Instead, in rare occurrences, additional pairs of +1 and −1 defects occur in axon patterning (Figure 1c). We also investigated cell shapes in the corneal epithelium directly (see Figure 3d), and found no evidence of systematic cell elongation or nematic order. We infer that here the polar, crawling motility of the cells dominates over cell elongation or stresses with nematic symmetry. Therefore, cell motion in a cornea is an example of a dense polar soft active matter system (***Marchetti et al., 2013***; ***Alert and Trepat, 2020***). This is the same conclusion (***Lång et al., 2024***) reached for HaCaT keratinocytes in a disk geometry but contrasts with the results of (***Guillamat et al., 2022***) for C2C12 myoblasts on a disk, where cells feature elongated shapes and are therefore primarily nematic.

**Figure 1.**
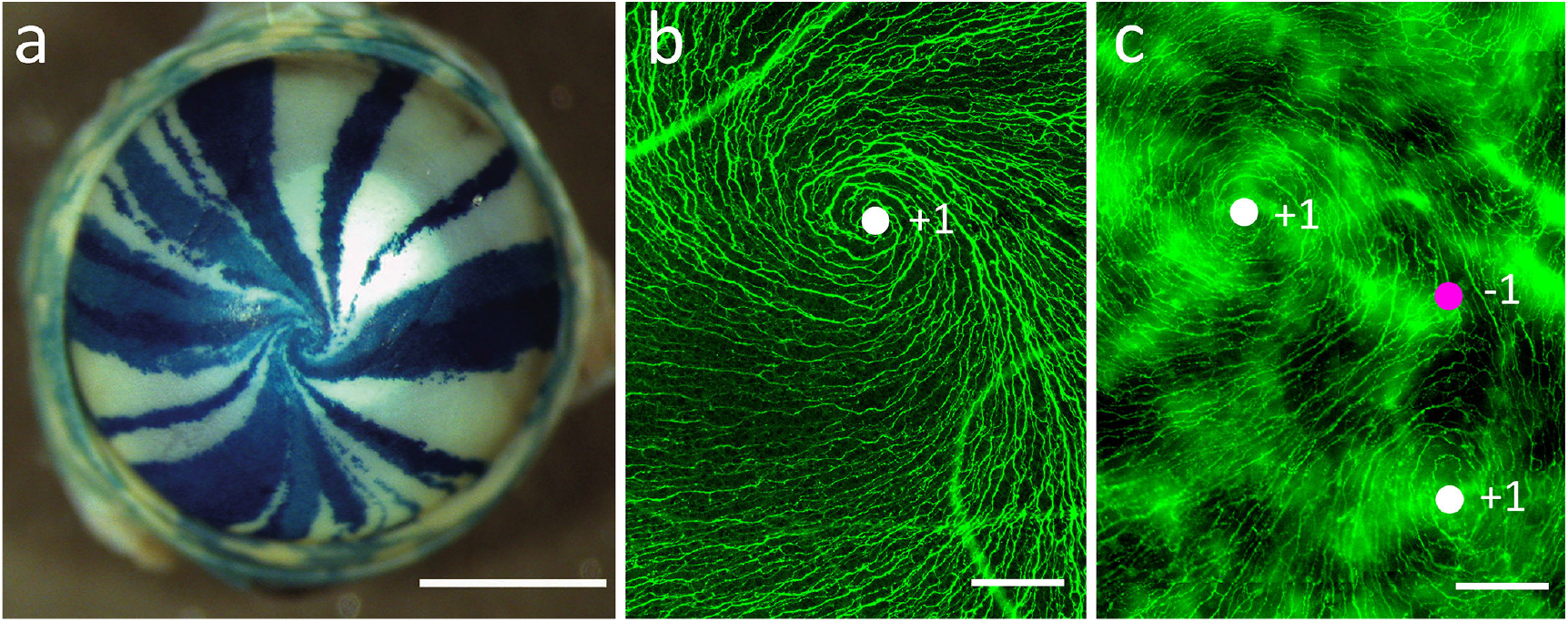
Spiral corneal striping of the murine eye. **a** LacZ-mosaic (‘XLacZ’) mouse cornea with radial spiral striping patterns of XGal staining, visualising patterns of epithelial cell migration during normal life. Image taken from (***Collinson et al., 2002***). **b** Beta-III-tubulin immunofluorescence staining of sensory axons in the spiral region of the basal layer of corneal epithelial cells. The axons follow the migratory flow of the basal epithelium. **c** A rare (≈1 in 100) occurrence of a double-swirl near the corneal centre. In both immunofluorescence images, integer topological defects (e.g. centres of swirls) can be readily seen. In **b** there is a single +1 defect (white circle). The unusual pattern, shown in **c**, shows two +1 defects and a single −1 defect (magenta circle). Scale bars: **a** 1000 µm. **b, c** 50 µm.

**Figure 2.**
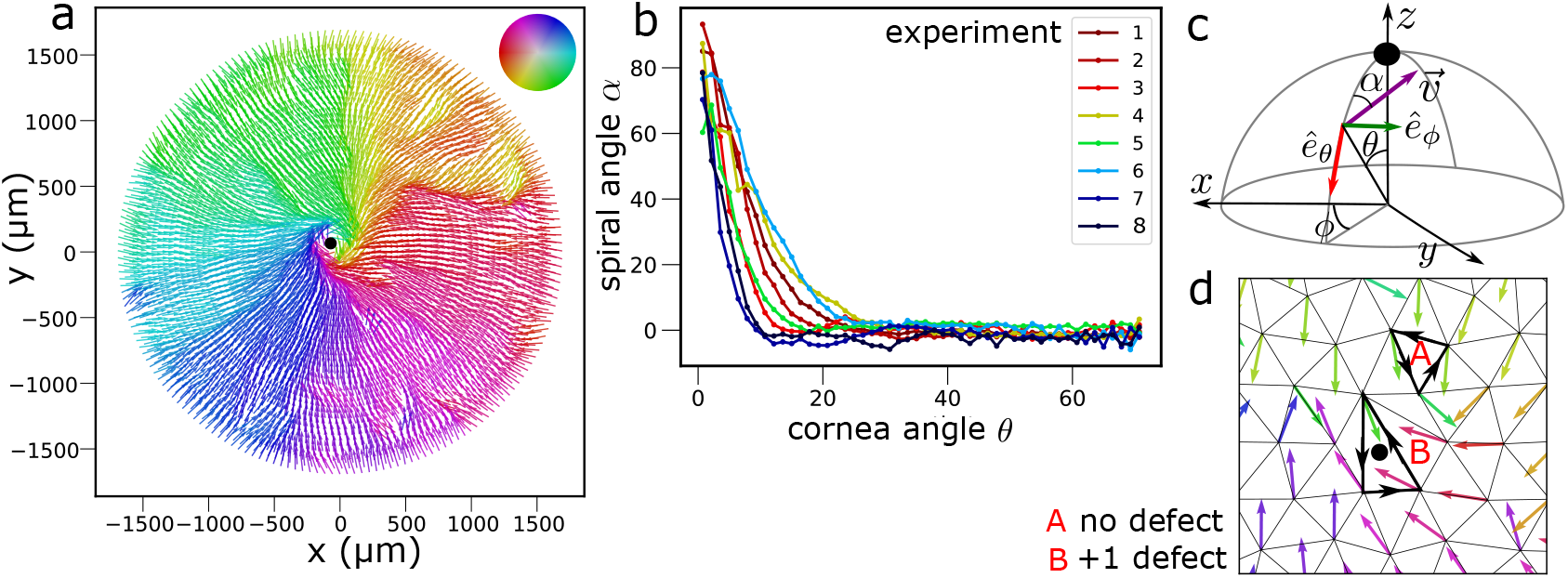
Experimentally observed corneal flow profiles. **a** Projection of the inferred flow field of experiment 1 onto the plane, where the colour corresponds to the velocity direction (see colourwheel legend). **b** Azimuthally (*ϕ*) averaged spiral angle *α* as a function of polar cornea angle *θ* from the central defect for *n* = 8 wild-type eyes (*θ* = 0 represents cornea center). **c** Local spherical coordinates around the location of the central topological defect (black). The angle *α* between the velocity vector and the −*ê*_*θ*_ direction defines the spiral angle as a function of the cornea polar angle *θ*. **d** Finding topological defects. From a Delaunay triangulation with the velocity arrows at the nodes, we sum up the angle differences Δ around each triangle and compute *c* = Δ/2*π*. These charges identify topological defects, e.g. *c* = 0 and no defect in the triangle labelled **A**, and *c* = 1 and a +1 defect in the triangle labelled **B**. **Figure 2–Figure supplement 1**. Fixed, stained and dissected LacZ-mosaic corneas used for flow inference. **Figure 2–Figure supplement 2**. Inferred flow fields for corneas 1-8.

**Figure 3.**
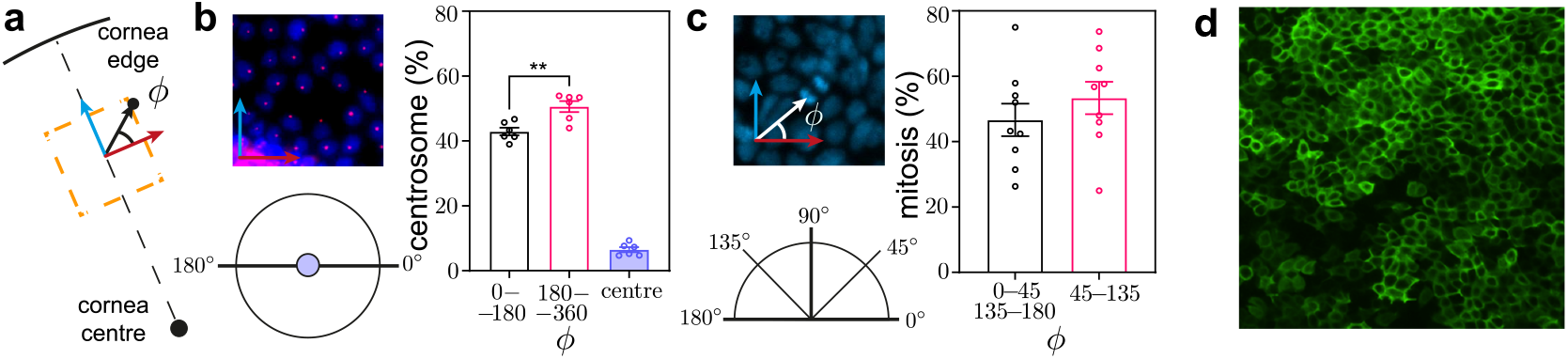
**a-c** Angles of centrosome position and mitotic spindle in the cornea during migration and cell division, respectively, showing no global angular correlations. **a** Definition of the division angle *ϕ* with respect to the corneal centre and edge. **b** Angles *ϕ* of centrosome position in relation to the nucleus and the centre of the cornea. The image shows nuclei stained with DAPI (blue), with the centrosome stained with pericentrin (pink). The position of the centrosome with respect to the nucleus may indicate an intracellular planar polarisation: our data suggested that centrosomes are slightly but significantly biased to lie in front of the nucleus *p* = 0.0047 (1-way ANOVA with Tukey’s posthoc multiple comparisons test). **c** Angles of mitotic spindle in relation to the centre of the cornea. The image shows a dividing cell with its mitotic spindle on a DAPI stained cornea. No preferential angle of division was observed. **d** Corneal epithelium flat mount immunostained for Keratin-12 intermediate filaments, highlighting the cell cytoplasm and boundaries. Cells show no preferred axis of elongation (i.e. nematic order).

To test the hypothesis of polar spiral migration, we reconstructed the epithelial cell flow fields across the corneal surface, by dissecting, flattening and imaging corneas as described in *Materials and Methods*. We manually traced boundaries between blue and white stripes from images of *n* = 8 dissected wild-type adult XLacZ corneas in Inkscape as described in the Materials and Methods (see Figure 11). We wrote custom analysis software in MATLAB to digitally reconstruct the cornea surface in 3D and then inferred the local cell flow directions from the stripe edges. Finally, we minimised an XY model of polar alignment to interpolate between known directions. This enabled us to infer the direction of velocity vectors everywhere on the cornea. In Figure 2a, we show a two-dimensional projection of the flow pattern along the curved corneal surface inferred from one of the corneas (experiment 1). For all corneas, striping patterns are shown in Figure 2 – Figure supplement 1, and inferred flow patterns are shown in Figure 2 – Figure supplement 2.

We identified topological defects in the flow field by computing angle differences in the projected velocity directions over loops defined by a Delaunay triangulation (***Zapotocky et al., 1995***; ***Henkes et al., 2018***) (Figure 2d). This method is a discretised version of the loop integral around a defect core shown in Box 1. We always found a single +1 topological defect surrounded by a spiral, located near the centre of the cornea (Figure 2a and Figure 2 – Figure Supplement 2).

We quantified the flow direction by the flow angle *α*, determined as follows. Owing to the nearly spherical shape of the eye, we used spherical coordinates to identify points on the cornea. We placed the origin at the centre of the eye, and the location of the +1 topological defect represents the ‘north pole’, such that the *z*−axis passes through it (Figure 2c). We measured the angular distance *θ* from the pole (like an inverse latitude), where *θ* = 0° corresponds to the spiral centre and *θ* ≈ 70° corresponds to the limbal region at the periphery of the mouse cornea. We measured the azimuthal direction by the angle *ϕ*, with values between 0° and 360°, with an arbitrary origin as the cornea is radially symmetric.

The spherical coordinates allowed us to define local unit reference vectors ê_*θ*_ and ê_*ϕ*_ in the tangent plane of the corneal surface. Then we defined *α* as the angle between − ê_*θ*_ and the direction of the velocity vector **v** (Figure 2c). Here the minus sign ensures that |*α*| *<* 90° for an inward flow, as ê_*θ*_ points outwards by convention.

The flow pattern is clearly highly symmetric along the azimuthal direction. Therefore, we averaged *α* over the angle *ϕ* to extract the characteristic spiral angle flow profile *α*(*θ*), as a function of the polar angle *θ* (Figure 2b). Although both clockwise and counterclockwise patterns were approximately equally likely, we flipped the sign of angle *α* on clockwise-rotating corneas for visual clarity. This enabled us to identify a reproducible pattern for all corneas: Flow is straight inward near the corneal edge with *α*(*θ* = 70°) ≈ 0 and stays such until at an angle *θ* ≈ 15° − 25°, when *α*(*θ*) sharply increases, corresponding to a systematic tilt of **v**, i.e. the onset of a spiral. In all cases, the spiral angle further continuously tightened until reaching *α*(*θ* = 0°) ≈ 90° at the corneal centre, corresponding to flow in the azimuthal direction. Below, we will show that this flow pattern, namely perpendicular anchoring and a vortex defect, is a natural consequence of the XYZ hypothesis.

### Excluding prepatterning as the stripe formation mechanisms

Patterns of radial striping are potentially explicable if the corneal epithelial cells are aligned by a global patterning mechanism that, e.g. biases cell migration or aligns cell division axes to point towards the cornea centre.

To test whether the cell migration direction is biased, we used the centrosome. The position of the centrosome has been associated with cell polarity, motility, and division (***Nigg and Raff, 2009***). Intracellular movement of the centrosome has been shown both in vivo and in wound assays in vitro (***Werner et al., 2017***; ***Blitzer et al., 2011***). Hence, the position of the centrosome relative to the nucleus can act as a proxy measurement for the cell polarity and, potentially, the direction of migration. To assay whether the corneal epithelial cells have an intrinsic directional polarity, we performed immunohistochemistry to visualise pericentrin in flat-mounted corneas and measured the angle of the centrosome-nuclear axis *ϕ* with respect to the corneal centre (6 eyes, 879 cells) (Figure 3a,b). Centrosomes were either behind the nucleus (0°−180°) (42±1%), in front of the nucleus (51 ± 2 %), or directly above or below the nucleus with no orientation (6 ± 1%). Statistically, there was a slight but significant tendency for centrosomes to locate in front of the nucleus (*p* = 0.0047), consistent with the observed mean flow towards the cornea centre. However, the overall scattered pattern of centrosomes with respect to the nucleus is more consistent with short-term changes in migration directions associated with the local cell environment than with any global patterning.

To investigate whether radially-aligned cell division axes contributed to observed striping patterns, we identified mitotic figures in DAPI-stained flat-mounted corneas from adult mice as shown in Figure 3. We manually measured the angle *ϕ* of each mitotic division with respect to the centre of the cornea and binned it as ‘radial division’ (0° − 45°, 135° − 180°) or ‘tangential division’ (45° − 135°). Figure 3c shows that we did not observe any significant difference in the proportions of radial (47 ± 5 %) and tangential (53 ± 5%) divisions (9 eyes, 376 dividing cells). Therefore, we did not find any evidence for radial striping being caused by biased axes of cell division.

### Soft active particle model of epithelial migration and parameter extraction

With the prepatterning mechanism being unlikely, we turn to describing the observed radial migration and spiral patterns as emergent phenomena. We therefore constructed an agent-based model rooted in the physics of soft active matter. To first establish a baseline before introducing curvature, we developed and parametrised a two-dimensional model using corneal explants and monolayers of primary mouse corneal epithelial cells.

The corneal epithelium is stratified into several layers, but active migration is performed solely by the basal layer of cells, while the top layers ride passively on the basal flow until desquamation. For simplicity, we therefore modelled the tissue as a single sheet of cells represented by soft disks, following overdamped Langevin dynamics. Each disk has a soft repulsive core of radius *σ*, accounting for elastic overlap between neighbours, and a short-range attractive potential modelling cell–cell adhesion. Active migration on the basement membrane is introduced through a self-propulsion force of magnitude *F*_act_ = *ζ υ*_0_ with direction determined by the unit vector **n̂**_*i*_ that reorients over time. Here, *υ*_0_ is the self-propulsion velocity, and *ζ >* 0 is the friction coefficient with the substrate. Cells can divide or be extruded with rates *d*_*z*_ and *a*, respectively. The division rate reduces with local density as *d*_*z*_ = *d*_0_(1 − *z*/*z*_max_), where *d*_0_ is the base rate and *z* is the number of neighbours of the cell, and is zero for *z* ≥ *z*_max_. Here, extrusion represents basal cell differentiation and upward movement into apical layers and *a* is taken as constant. Together, these allow for homeostasis when division balances extrusion, with a typical cell division time of *a*^−1^ (***Matoz-Fernandez et al., 2017***).

This simple model of epithelial cell migration, previously used to model human corneal cell layers (***Henkes et al., 2020***), is of the Active Brownian class of active materials (***Marchetti et al., 2016***) while motility directions 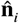 remain uncoordinated between cells. Its collective cell migration patterns are ‘swirly’, with mesoscale spatiotemporal correlations set by the interaction of cell elasticity with persistence time of migration (***Henkes et al., 2020***; ***Kammeraat et al., 2025***) matching those seen, e.g. in MDCK monolayers (***Angelini et al., 2011***). It has been recognised (***Keta et al., 2024***) as one of several mechanisms that generate active ‘turbulence’ - a collective, disordered motion arising from internally generated forces produced by cells themselves - in cell sheets and other systems like bacterial liquids (***Alert et al., 2022***). This simple model, however, does not allow for large-scale collective cell migration, i.e. flocking, at the scale of the cornea.

Cells also coordinate their migration direction through local mechanical or chemical cues. Although the detailed mechanochemistry of corneal epithelia remains unknown, studies of MDCK monolayers have identified multiple coordination modes, including plithotaxis—reorientation along principal stress directions (***Trepat and Fredberg, 2011***). In the absence of force traction or myosin expression data, we included two reorientation mechanisms: (i) alignment with the local force on the cell, i.e. internal force feedback (***Szabó et al., 2006***; ***Henkes et al., 2011***; ***Petrolli et al., 2019***; ***Malinverno et al., 2017***; ***Peyret et al., 2019***; ***Lång et al., 2024***), and (ii) alignment with the migration direction of nearest cell neighbours (***Saraswathibhatla et al., 2021***). Both mechanisms can generate collective migration (‘flocking’) and belong to the class of polar active materials (***Giardina, 2008***; ***Vicsek and Zafeiris, 2012***; ***Baconnier et al., 2025***).

Our model assumes that the dominant source of motility is from cells crawling on the substrate, as opposed to cell-cell stresses transmitted through junctions or coming from cell elongation (***Alert and Trepat, 2020***). The latter assumption leads to *active nematic* models of cell motility (***Doostmohammadi et al., 2018***; ***Shankar et al., 2017***; ***Duclos et al., 2018***), leading to a prominent role of half-integer ±1/2 defects (***Saw et al., 2017***; ***Yu et al., 2025***). As detailed above, we adopted a polar instead of nematic active particle model because we observed neither systematic cell elongation (Figure 3d) nor ±1/2 topological defects (see Box 1 Figure 1c-d) in the axon patterns in Figure 1b-c. Thus, each cell can generate motion individually, and the model captures collective behaviour through polar alignment, leading to ±1 defects. We emphasise that ‘polar’ here refers to the physical symmetry of the active forces, not to biological cell polarity.

Ultimately, the equations of motion for cell position **r**_*i*_ and migration angle *ϑ*_*i*_ in our model are:

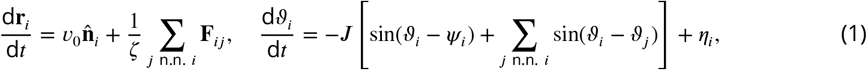

where the migration direction is written as 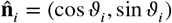, with *ϑ*_*i*_ the angle of 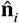 and *ψ*_*i*_ the angle of the cell velocity (or force) vector **v**_*i*_, both measured from the *x*−axis of the simulation box. The mechanical forces **F**_*ij*_ derive from a short-range potential that includes repulsion and attraction with stiffness *k, J* is the alignment strength, and *η*_*i*_ is a rotational white noise with ⟨*η*_*i*_(*t*)*η*_*j*_(*t*′)⟩ = 2*D*_*r*_*δ*_*ij*_*δ*(*t* − *t*′), defining the persistence time *τ* = 1/*D*_*r*_. Details of the numerical implementation are provided in *Materials and Methods* and (***Sknepnek and Henkes, 2015***; ***Henkes et al., 2020***); we used our simulation package (***SAMoS, 2024***) to carry out the computations and our matching analysis package (***SAMoSA, 2024***) to analyse the data.

To parametrise the model, we measured cell sizes, velocities, flow patterns, and cell cycle rates in mouse corneal epithelial explants (‘explant’) where the basal epithelial cells move on their own stromal layer and monolayers cultured on plastic substrates (‘plastic’) (Figure 4). The mean basal cell radius as directly measured from microscopy was *σ* ≈ 5 − 6 µm in explants and *σ* ≈ 11 µm on plastic. The mean cell cycle time in mouse basal corneal epithelial cells has been measured previously to be ≈ 3.5 − 5.0 days (***Sagga et al., 2018***), with individual basal epithelial cells capable of division once every 2.5 days (***Lehrer et al., 1998***).

**Figure 4.**
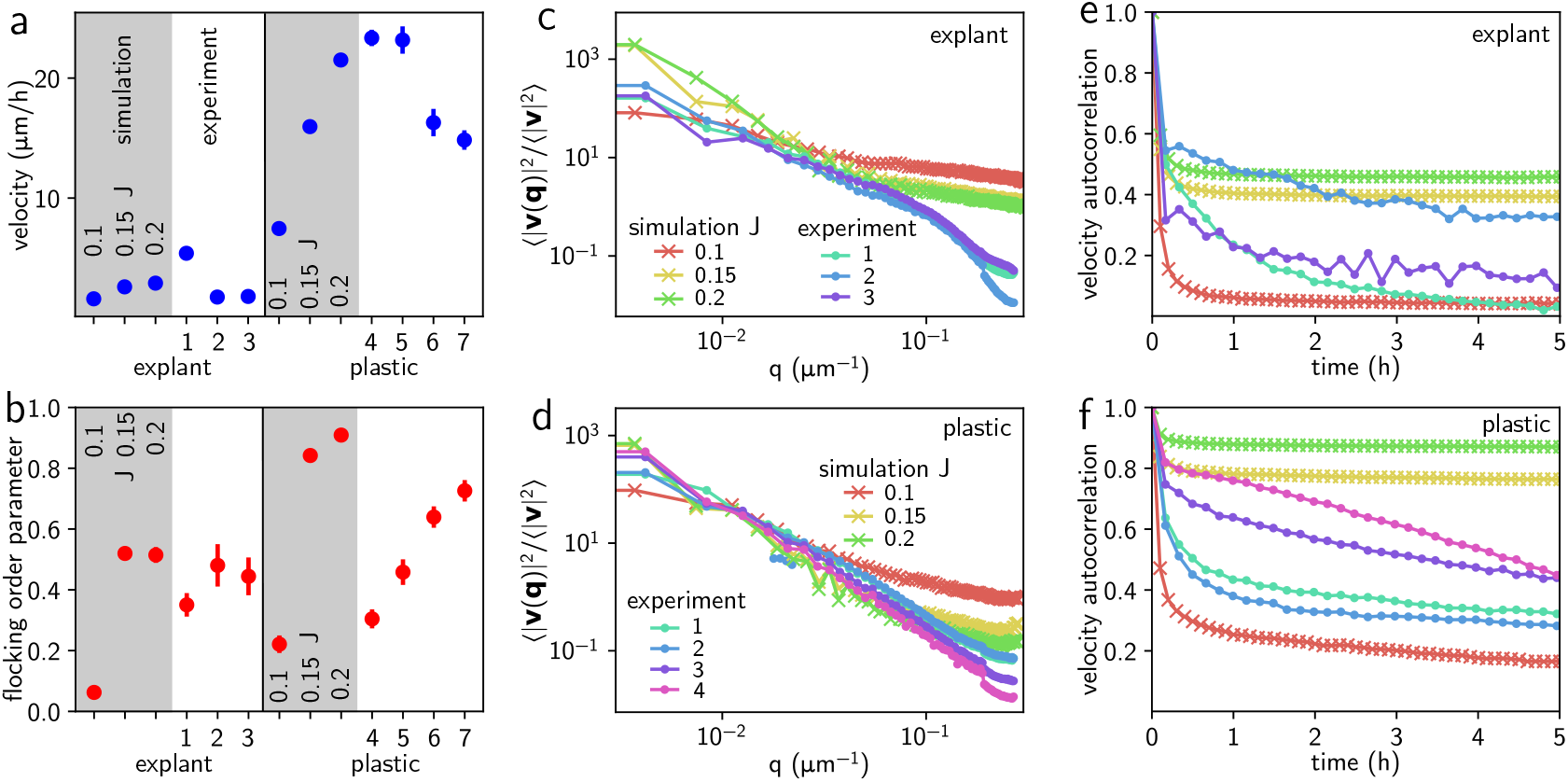
Matching procedure of simulated cell flows to experiment, for corneal-derived cells on the stromal substrate within the explant (labelled ‘explant’, *n* = 3), and for cells proliferating on tissue culture plastic substrate (labelled ‘plastic’, *n* = 4). For both conditions, we included 3 simulations with all parameters fixed except for the alignment strength, *J*, and one simulation with additional pair friction (labelled ‘f’). **a** Root mean squared cell velocity magnitude 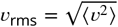 in simulations (shaded regions) and for experiments (white regions). Error bars represent one standard deviation. If not shown, the error bar is smaller than the symbol size. **b** Flocking order parameter *ϕ* = |⟨**v**⟩| /*υ*_rms_, with shading and symbols same as in **a. c** and **d** Normalised velocity correlation function ⟨**v**(*r*) ⋅ **v**(0) ⟩ / ⟨*υ*^2^⟩ for the ‘explant’ and ‘plastic’ conditions, simulation curves are for the hydrodynamic velocity (see text). **e** and **f** Normalised velocity autocorrelation function ⟨**v**(*t*) ⋅ **v**(0) ⟩ / ⟨*υ*^2^⟩ for ‘explant’ and ‘plastic’ conditions. **g** Corneal explant overlaid with velocity arrows from PIV analysis and **h** simulated explant at *J* = 0.15. **i** Cells on the plastic substrate overlaid with velocity arrows and **j** simulated plastic substrate at *J* = 0.15. Arrows for the simulations are the hydrodynamic velocity.

Cell motility patterns, as determined by PIV analysis, showed signs of both ‘swirly’ and ‘flocking’ motility, see Figure 4 g,i and supplementary videos S1 and S2. We characterised them using the following measures: First the root-mean-square velocity 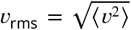 (where ⟨⋅⟩ denotes an average taken over all cells and over time), ranging from 2–6 µm h^−1^ in explants to 15–25 µm h^−1^ on plastic (Figure 4a), consistent with higher traction forces on stiffer substrates (***Ladoux and Mège, 2017***), and in a similar region as previous experiments (***Leiper et al., 2006***; ***Garcia et al., 2015***; ***Walczysko et al., 2016***). Second, the flocking or Vicsek order parameter Φ_*υ*_ =|⟨ **v** ⟩| /*υ*_rms_, a number that ranges from 0 (no collective motion) to 1 (perfect flock). Experiments on both substrates fall into the 0.3 − 0.7 range (Figure 4b), indicating that our epithelia are right around the threshold of the flocking phase transition (***Vicsek et al., 1995***). Third, we analysed the spatio-temporal correlations of the velocity field. Specifically, we measured (i) the spatial velocity correlation function, ⟨**v**(*r*) ⋅ **v**(0) ⟩ / ⟨*υ*^2^⟩, which quantifies how similarly cells move as a function of their separation, and (ii) the temporal velocity correlation function, ⟨**v**(*t*) ⋅ **v**(0) ⟩ / ⟨*υ*^2^ ⟩, which measures how persistent cell motion is over time. In both cases, the mean collective (flocking) velocity was subtracted before computing the correlations. The spatial correlations are characterised by a correlation length of approximately *ξ* = 150*µm* for the plastic substrates, and a shorter initial decay *ξ* = 50*µm* for the corneal explants, but with a long tail. The temporal correlations are all characterised by an initial rapid decay followed by a long tail or plateau, with the correlations on the plastic substrate again being stronger.

We matched these results using our model with the following parameters: We used the measured cell radii *σ* = 5µm in explants and *σ* = 11 µm on plastic, and the mean cell cycle set the extrusion rate to *a*^−1^ = 50 h. Then the division rate *d*_*z*_ was tuned to achieve steady-state density above confluence (***Matoz-Fernandez et al., 2017***).

Corresponding self-propulsion speeds were *υ*_0_ = 3 µm h^−1^ and 30 µm h^−1^, allowing for *υ*_rms_ values in the range of 2–6 µm h^−1^ in explant simulations and 15–25 µm h^−1^ on plastic (Figure 4a, grey regions labelled ‘simulations’). The alignment strength *J* was estimated by matching both *υ*_rms_ and the flocking order parameter Φ_*υ*_ = |⟨ **v** ⟩| /*υ* _rms_: values in the range *J* = 0.1 − 0.2 h^−1^ matched experimental data (Figure 4b) and correspond to the onset of collective migration. The ratio *k*/*ζ*, which determines the elastic response time, was adjusted to 30 h^−1^ to reproduce the observed velocity correlations in space (Figure 4c,d) and the persistence time *τ* = 2h to match the velocity correlations in time (Figure 4e,f), similar to the procedure detailed in (***Kammeraat et al., 2025***).

In addition to the semi-quantitative matches, with simulations that look appropriately swirly and flocking (see Figure 4h, j for snapshots and supplementary videos S3 and S4), we also observe the characteristic features previously noted: For the spatial correlations, shorter correlations are followed by a tail, and the appearance of a plateau after a rapid decay in the temporal correlations. Both features increase with alignment strength *J*, pointing to them arising from the combination of swirling and flocking motility.

We note that the way cell division and extrusion are implemented in the model generates locally opposing velocity pairs, leading to negative velocity correlations at short ranges (see Figure 4 d,h). We therefore used a time-averaged, or hydrodynamic cell velocity from cell displacements 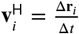, excluding divisions. There has also been recent insight that active flow correlations sensitively depend on the presence or absence of a pair friction term, *ζ*_*f*_ (**v**_*i*_ − **v**_*j*_), with correlations decaying less abruptly in its presence (***Tong et al., 2023***; ***Rozman et al., 2025***; ***Kammeraat et al., 2025***). We therefore also simulated the model with a *ζ*_*f*_ = *ζ* = 0.5 instead of the usual *ζ* = 1, *ζ*_*f*_ = 0. We do find results for the hydrodynamic velocity that are comparable to each other, albeit at lower values of alignment *J*.

Overall, the fitted parameters (Table 1) produced quantitatively accurate velocity statistics for both explant and plastic conditions (Figure 4g–j; Videos S1–S4), validating the model as a faithful representation of collective migration in corneal epithelial cells.

**Table 1.** Model parameters and their values inferred from experiment and used in simulations. Bare values for the cell-cell interaction potential strength *k* and the friction coefficient *ζ* are not shown since only their ratio *k*/*ζ* matters in our model.

| Parameter | Description | Value |
| --- | --- | --- |
| $k/\zeta$ | soft potential strength | $30 \text{ h}^{-1}$ |
| $\epsilon$ | attraction depth and width | 0.15 |
| $\sigma$ | cell radius | $5 \text{ }\mu\text{m}$ ('explant'), $11 \text{ }\mu\text{m}$ ('plastic') |
| $J$ | alignment strength | $0-0.2 \text{ h}^{-1}$ |
| $D_r$ | rotational diffusion constant | $0.5 \text{ h}^{-1}$ |
| $v_0$ | self-propulsion velocity | $3 \text{ }\mu\text{m/h}$ ('explant'), $30 \text{ }\mu\text{m/h}$ ('plastic') |
| $a$ | cell extrusion rate | $0.02 \text{ h}^{-1}$ |
| $d_0$ | base rate of cell division | $0.2 \text{ h}^{-1}$ |
| $z_{\max}$ | maximum number of neighbours | 8 |
| $R$ | cornea radius | $100 - 1500 \text{ }\mu\text{m}$ |
| $dt$ | time step | $0.001 \text{ h}$ |

**Table 2.** Key resources table.

| Reagent type (species) or resource | Designation | Source or reference | Identifiers | Additional information |
| --- | --- | --- | --- | --- |
| strain, strain background (Mus musculus) | XLacZ mouse; H253 | Collinson et al., 2002 | X-linked LacZ transgene | Adult female mosaic mice used for corneal striping analysis |
| biological sample (Mus musculus) | H253; Adult mouse corneas | This paper |  | Flat-mounted, dissected and X-Gal stained corneas for flow-field inference |
| cell line (Mus musculus) | Primary mouse corneal epithelial cells | Henkes et al., 2020 |  | Cultured on tissue culture plastic and corneal explants |
| antibody | anti-pericentrin | Abcam | Ab448 | Immunohistochemistry for centrosome localisation |
| stain | DAPI | Vector labs | H-1200 | Nuclear staining of corneal epithelial cells and mitotic figures |
| software, algorithm | Inkscape | Inkscape Project |  | Manual tracing of stripe boundaries |
| software, algorithm | MATLAB | MathWorks |  | Custom analysis software for flow-field reconstruction |
| software, algorithm | SAMoS | R. Sknepnek, https://github.com/-sknepneklab/SAMoS |  | Simulation package used for active particle simulations |
| software, algorithm | SAMoSA | S. Henkes, https://github.com/-silkehenkes/SAMoSA |  | Analysis package used for active particle simulations |

### Corneal surface model

To combine active cell dynamics with the realistic three-dimensional corneal shape, we modelled the cornea as a spherical cap of radius *R* = 100 − 1500 µm and a 70° opening, consistent with experimental measurements. The corneal epithelial cells, i.e. the TA cells derived from stem cell divisions in the limbus, were modelled as soft particles able to migrate on the curved surface, divide, and be removed. To ensure that the cells remain confined to the surface throughout the simulation, we projected the velocity vectors onto the local tangent planes at each time step. The equations of motion, therefore, contain the same term as in the flat case, eqn. (1), with additional projection operators (***Sknepnek and Henkes, 2015***) (see *Materials and Methods*). The model parameters, listed as ‘explant’ in Table 1, were determined as discussed in the previous section. At the bottom boundary, we modelled the limbal stem cell niche as a single layer of continuously dividing cells, a simplification that takes into account both dividing stem cells and a population of TA cells that may divide before leaving the limbus (***Vanin et al., 2025***).

In the mouse limbal epithelium, slow-cycling stem cells, comprising approximately 25% of the cell population, were previously determined to divide once every 10 - 20 days with the other 75% of rapidly dividing cells having a cell cycle time of 3 - 3.5 days (***Sagga et al., 2018***). To match the above and the empirically determined corneal resurfacing time of 14 days at full-size, we chose a density-independent division rate of *d*_*L*_ = 0.4 h^−1^ for the limbal ring.

Finally, we included a double ring of tightly spaced stiff cells as a barrier to constrain motion to the interior of the cornea area. This reflects experimental observations that the progeny of limbal epithelial stem cells never migrate outwards into the conjunctiva. We recapitulated the lineage tracing provided by XLacZ staining by separating the cells into ‘blue’ and ‘white’ types. Each ‘blue’ limbal stem cell gives rise to ‘blue’ TA cells, which can also only divide further into more ‘blue’ cells, as in vivo. We used the same dynamics for the ‘white’ cells. Blue and white cells were identical in their interactions and they could mix freely. We set the base for the stripes to 12 alternating regions of fixed stem cells along the limbal stem cell line (Figure 5). For the initial set-up of simulations, we randomly seeded cells with a 50:50 ‘blue’:’white’ mixture on the corneal surface. For the largest cornea radii *R*_c_ = 1000 µm and *R*_c_ = 1500 µm, due to computational limitations, we started with an empty corneal surface. As above, we implemented the model through custom configuration files and initialisation scripts in our package (***SAMoS, 2024***). We simulated between 1400 cells at *R*_c_ = 100 µm, and 241 000 cells at *R*_c_ = 1500 µm. We ran the simulations using an Euler-Maruyama algorithm for 500 000 simulation time steps at d*t* = 0.001 h, corresponding to 20.8 days of development. Please see supplementary video S5 for a *R* = 100 simulation corresponding to Figure 5.

**Figure 5.**
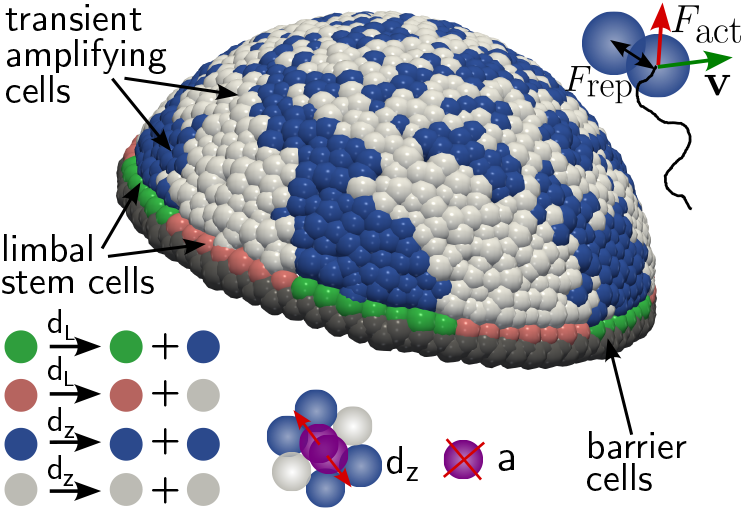
In silico corneal model. Cells were constrained to move on the surface of a spherical cap with a polar angle of 70°. Each cell is self-propelled by an internal active force, *F* ^act^, in addition to being subject to a short-range elastic force *F*_el_ due to mechanical interactions with neighbouring cells. Due to slow migration speeds, cell motion is overdamped, and forces are balanced by friction with the substrate, giving rise to cell velocity **v.** Inwards from a layer of immobile barrier cells (grey), the stem cells of the limbal niche (green and pink) divide with rate *d*_*L*_ to give rise to transient amplifying (TA) cells (blue and white). TA cells divide with the density-dependent rate *d*_*z*_ into cells with the same label and are extruded with rate *a*.

### Emergence of the spiral pattern

The corneal spiral is not present from birth (Figure 6a): Young XLacZ mice exhibit a random patchwork arrangement of blue and white clones from mosaic LacZ expression of cells. Cell loss stimulates the immigration of cells into the cornea from the limbal epithelium, such that by adulthood the patterns resolve as the spiral radial arrangement of clones. The transition from a primarily patchwork to a primarily striped pattern takes 3 - 4 weeks, with intermediate patterns with stripes emerging from the periphery, retaining a patchwork centre. We were able to successfully reproduce both this transition and the spiral striping pattern using numerical simulations of the corneal surface model. In Figure 6b-e, we show the time evolution of a simulated cornea of radius *R* = 700 µm. As can be seen in the second row (Figure 6b and accompanying supplementary video S6), the initially randomly distributed ‘blue’ and ‘white’ cell pattern was gradually replaced by a striped pattern made of cells derived from the limbus. This was not simply a case of cells being replaced since the change happened over several generations of cells, with the overwhelming majority of striping pattern cells being derived from other corneal epithelial TA cells. By day 20, a cornea with a stable central spiral emerged. This pattern of replacement, including the time scale over which it occurred, was very similar to the corneal evolution observed in neonatal mice shown in Figure 6a. We can understand the emergence of the spiral flow pattern by examining the temporally averaged (or hydrodynamic) velocity field (Figure 6c and supplementary video S7), where we tracked the displacement Δ**r**_*i*_(*t*) = **r**_*i*_(*t* + Δ/2) − **r**_*i*_(*t* − Δ/2) of each surviving cell *i* over Δ*t* = 2.5 h, giving 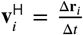. This is distinct from the instantaneous cell migration speed **v**_*i*_ which was more random, consistent with the nearly random cell orientations seen in experiments (cf. Figure 3b). Figure 12 in the *Materials and Methods* shows how the smooth hydrodynamic flow pattern emerges as we increase the averaging time Δ*t*. The hydrodynamic velocity field gradually forms a spiral pattern with flow parallel to the blue/white stripe edges of the spiral. This confirmed our experimental approach of inferring the flow field direction from the stripe edges.

**Figure 6.**
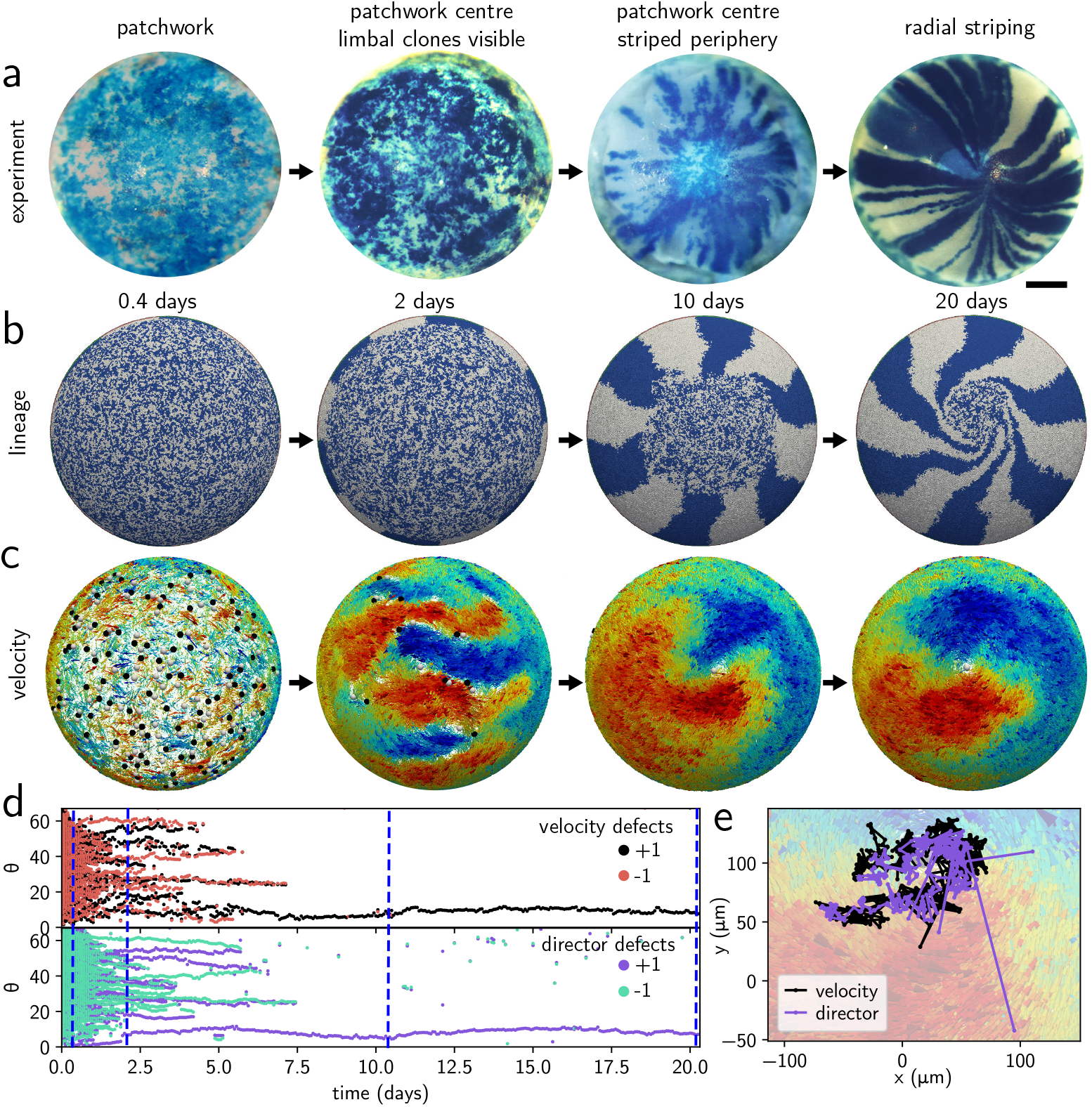
Emergence of the spiral flow in experiment and in a simulated *R* = 700 µm cornea. **a** X-Gal-stained beta-galatosidase mosaic corneas from ‘H253’ (XLacZ) female mice, hemizygous for X-linked LacZ transgene (***Collinson et al., 2002***), showing the gradual appearance of the spiral over 3 - 4 weeks. Scale bar is 500 µm. **b** Simulated time sequence of the emerging spiral pattern with labelled ‘blue’ and ‘white’ TA cells. The spiral is completely formed by simulated day 20. **c** Corresponding hydrodynamic flow fields **v**^H^ of the cells, coloured according to the *x* (left-right) component of the flow direction in 3D space from blue to red. The flow is determined by topological defects (white +1, black −1) that merge to give rise to a central +1 topological defect by day 8. **d** Velocity (top) and director (bottom) topological defect polar angle as a function of time. +1 and −1 defect pairs merge progressively, the snapshots in **b** and **c** correspond to the blue vertical lines. **e** Central +1 defect track for both velocity and director overlaid on the velocity field.

We next quantified our observations by tracking the +1 and −1 topological defects of **v**^H^, shown in white and black, respectively in Figure 6c. As discussed in Box 1, topological defects correspond to regions where the flow field is discontinuous, with +1 defects being circular objects with spiral flows around the defect core, while −1 defects have four-fold inward/outward symmetry (see Box 1 Figure 1a-b). We computed topological defect charges *c* using the same discrete sum around Delaunay triangles of a tesselation constructed from the particle positions as for the experiment (Figure 2d), and further merged the charges of defects recursively within 15 µm, to avoid regions of small-scale disordered flow dominating our measurements (***Henkes et al., 2018***; ***Prathyusha et al., 2018***). Please see *Materials and Methods* for full details; both the computation of **v**^H^ and the topological defects were carried out in our analysis package (***SAMoSA, 2024***). Since the cornea is topologically a disk, i.e. a surface with a boundary, in general, the total topological charge is influenced by that boundary. However, due to flows being uniformly inward close to the limbal region, corresponding to perpendicular anchoring, the sum of the topological charges is expected to be +1, i.e. *N*_+1_ − *N*_−1_ = 1 (***Stein, 1979***). We indeed observed that as the simulated cornea matures, +1 and −1 defects were initially very numerous and then reduced until only one central, +1 topological spiral defect remained, which corresponded to the centre of the blue-white spiral. In Figure 6d, we show the polar angle of the ±1 defects as a function of time. One can clearly see the mutual annihilation of +1 and −1 defects such that by day 8, only the central +1 defect remained. Here the top graph corresponds to the defects of the flow field shown in Figure 6b, and the bottom graph corresponds to the director field, i.e. the field of directions of migration of individual cells. Figure 6e shows the spatial trajectory of the central defect superimposed on the flow pattern, again for velocity and director defects. Both show that the central defect is almost stationary and nearly identical for velocity and director defects. These measures all show that the flow field pattern is a direct result of the polar alignment of the cells, i.e. the spiral pattern indeed emerges from the interaction between flocking and hemispherical geometry. We note that a similar pattern of defects annihilating in pairs into a central vortex was seen in (***Lång et al., 2024***) for both cells and a polar active model on a disk topology.We next quantified our observations by tracking the +1 and −1 topological defects of **v**^H^, shown in white and black, respectively in Figure 6c. As discussed in Box 1, topological defects correspond to regions where the flow field is discontinuous, with +1 defects being circular objects with spiral flows around the defect core, while −1 defects have four-fold inward/outward symmetry (see Box 1 Figure 1a-b). We computed topological defect charges *c* using the same discrete sum around Delaunay triangles of a tesselation constructed from the particle positions as for the experiment (Figure 2d), and further merged the charges of defects recursively within 15 µm, to avoid regions of small-scale disordered flow dominating our measurements (***Henkes et al., 2018***; ***Prathyusha et al., 2018***).

Please see *Materials and Methods* for full details; both the computation of **v**^H^ and the topological defects were carried out in our analysis package (***SAMoSA, 2024***). Since the cornea is topologically a disk, i.e. a surface with a boundary, in general, the total topological charge is influenced by that boundary. However, due to flows being uniformly inward close to the limbal region, corresponding to perpendicular anchoring, the sum of the topological charges is expected to be +1, i.e. *N*_+1_−*N*_−1_ = 1 (***Stein, 1979***). We indeed observed that as the simulated cornea matures, +1 and −1 defects were initially very numerous and then reduced until only one central, +1 topological spiral defect remained, which corresponded to the centre of the blue-white spiral. In Figure 6d, we show the polar angle of the ±1 defects as a function of time. One can clearly see the mutual annihilation of +1 and −1 defects such that by day 8, only the central +1 defect remained. Here the top graph corresponds to the defects of the flow field shown in Figure 6b, and the bottom graph corresponds to the director field, i.e. the field of directions of migration of individual cells. Figure 6e shows the spatial trajectory of the central defect superimposed on the flow pattern, again for velocity and director defects. Both show that the central defect is almost stationary and nearly identical for velocity and director defects. These measures all show that the flow field pattern is a direct result of the polar alignment of the cells, i.e. the spiral pattern indeed emerges from the interaction between flocking and hemispherical geometry. We note that a similar pattern of defects annihilating in pairs into a central vortex was seen in (***Lång et al., 2024***) for both cells and a polar active model on a disk topology.

### Key parameters of simulated spiral corneas

We next sought to understand the properties of the spiral flow field and how it compared to the experimental observations. In Figure 7, we show the dependence of the simulated flow patterns on cornea radius *R* and polar alignment strength *J*. Figure 7 – Figure supplement 1 and 2, respectively, show simulation snapshots of all simulated values of *J* and *R*.

**Figure 7.**
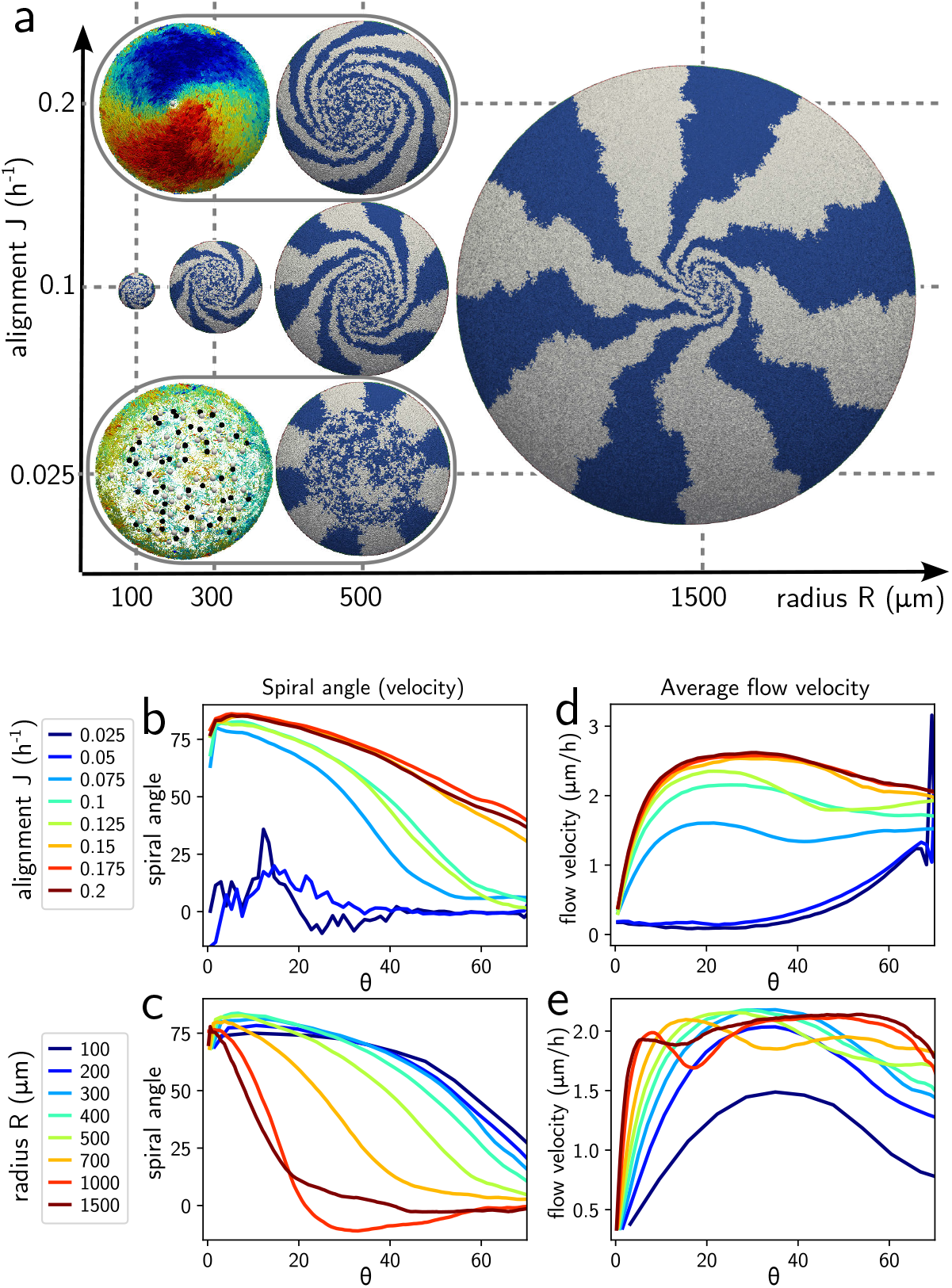
Properties of the spiral as a function of the corneal radius *R* and alignment strength *J*. **a** Schematic phase diagram with lineage tracing and select flow fields after 20 days. Spirals appear at all *R*, but need *J* ≈ 0.075 h^−1^ to emerge. We include our best fit for *R* = 1500 µm life-size cornea. **b** and **c** Spiral angle *α* with respect to the inward −ê_*θ*_ direction as a function of polar angle *θ* as a function of *J* for *R* = 500 µm (**b**) and as a function of *R* for *J* = 0.1 h^−1^ (**c**). **d** and **e** Coarse-grained flow velocity magnitude *υ* as a function of polar angle *θ* as a function of *J* for *R* = 500 µm (**d**) and as a function of *R* for *J* = 0.1 h^−1^ (**e**). **Figure 7–Figure supplement 1**. Emergence of the corneal spiral as a function of *J* for *R* = 500 µm simulated corneas, snapshots are at 20 days. **Figure 7–Figure supplement 2**. Emergence of the corneal spiral as a function of *R* for *J* = 0.1 h^−1^ simulated corneas, snapshots are at 20 days. Note that simulations for *R* = 200, 400 and 1000 µm have patterned 48 instead of 12 corneal stripes into the limbus.

From these simulations, we made a key observation: polar alignment is necessary for spiral formation, and the cornea is not effectively resurfaced in its absence. For example, in the bottom grey oval in Figure 7a, we show the flow field (left) and a snapshot of the cell configuration (right) for *J* = 0.025 h^−1^ and *R* = 500 µm after 20 days. The flow field remained disordered with numerous topological defects and no emerging spiral. Supplementary videos S8 and S9 show the time evolution (or lack thereof) of this process. Conversely, for *J* = 0.2 h^−1^ (top grey oval) we obtained a tighter spiral with a well-developed spiral flow field. Supplementary videos S10 and S11 show the time evolution, including the more rapid appearance of the single central defect.

We quantified these observations by computing azimuthally averaged profiles using spherical coordinates with the central defect along the *z* axis, using the same method as for the experiment (Figure 2c). In Figures 7b and 7d we, respectively, show the spiral angle *α* and the mean flow velocity *υ*^*H*^ as a function of polar angle *θ*. The former is the same measure as in Figure 2b obtained in experiments. We found a transition between a disordered state with small *υ*^*H*^ in the interior (and thus no effective resurfacing) and no developed spiral for *J* ≤ 0.05 h^−1^ and increasingly effective resurfacing for *J* ≥ 0.075 h^−1^, in the same region as the flocking transition in the flat case. Both spiral shape and flow velocity saturated for *J* ≈ 0.2 h^−1^, in the limit where cells fully flock together. Thus, we predict that the emergence of the topological spiral flow due to polar alignment is essential for a healthy cornea.

We observed that a spiral emerges for all simulated cornea radii (*R* = 100 − 1500 µm), showing the robustness of the flocking mechanism. However, the spiral shape is strongly radius-dependent. In Figure 7c, we show the spiral angle *α* with the inward direction as a function of polar angle *θ* for different values of *R*. We see that there is a systematic change from a pattern of a wide spiral (large *α* at most *θ*) at small values of *R* to a flux that is nearly straight inwards on the outer areas of the cornea (small *α* at large *θ*) followed by an increasingly sharper spiral pattern near the centre. In all cases, we had *α*(0) → 90°, i.e. a spiral that becomes a vortex at the centre. The flow field *υ*^*H*^ (Figure 7e) shows that all corneas except the smallest reached a steady flow velocity of approximately 2 µm/h on the corneal surface. Near the centre, however, *υ*^*H*^ drops to 0 approximately linearly, with an increasingly sharper gradient as the corneal size increases.

### The XYZ model revisited

A mainstay of understanding of ocular surface maintenance has been the XYZ hypothesis which states that the amount of cells flowing into the cornea from the limbal stem cell reservoir is exactly balanced by the difference between the amounts of TA cells being born and leaving the cornea (***Thoft and Friend, 1983***). In other words, the XYZ hypothesis posits that the cornea is in a steady state with a constant but nonzero flux of cells from the limbus inward to the tip of the cornea. We can test and expand on this hypothesis by writing a continuum conservation law for cell numbers, and apply it to the spiral flow profiles, linking cell flux ***υ***^*H*^, the spiral angle of the cornea *α*, and the difference between birth and extrusion rates.

The XYZ hypothesis states that the number of cells in a patch of the cornea is, on average, constant. If we denote the number of cells in a patch as *N*, *N*_in_ cells flowing into the patch, *N*_out_ cells flowing out of the patch, *N*_d_ cells dividing, and *N*_a_ cells that are extruded, we obtain the following balance relation:

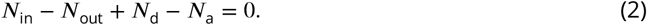

This equation could be reformulated in terms of the density of cells, *ρ* = *N* /*S*, where *S* is the patch area. The flux **J** = *ρ***v**^*H*^ is the mean number of cells with velocity **v**^*H*^ flowing through the boundary of *S* per unit time. Similarly, we can define *d*(*ρ*) and *a*(*ρ*), the division and death rates per cell. Eqn. (2) can then be reformulated as a flux conservation or continuity equation (see *Materials and Methods*) that states that the divergence of the flux at each point needs to be balanced by the difference in cells being extruded and those that are dividing at the same point, i.e.

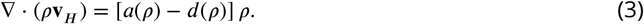

It is possible to link this equation with the XYZ hypothesis as follows. We first assume that the flow has a radially symmetric spiral pattern, i.e. **v**^*H*^ = *υ*(*θ*) cos *α*(*θ*)**e**_*θ*_+sin *α*(*θ*)**e**_*ϕ*_, where *υ*(*θ*) = |**v**^*H*^| is its magnitude and *α*(*θ*) is the angle between **v**_*H*_ and −**e**_*θ*_. The difference between division and extrusion can be combined into the density-dependent net loss rate *A*(*ρ*) = [*a*(*ρ*) − *d*(*ρ*)]. We further assume that the deviations of *ρ* from its mean are small, though we retain their influence on *A*(*θ*) ≡ *A*(*ρ*(*θ*)). Written in spherical coordinates for a cornea of radius *R* with the defect along *z* again as in Figure 2d, eqn. (3) becomes (see *Materials and Methods*),

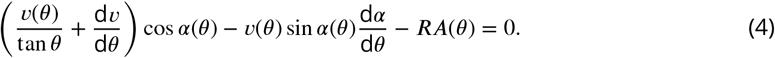

This equation links the spiral angle profile *α*(*θ*), shown in Figures 7b and 7c with its velocity profile *υ*(*θ*) (Figures 7d and 7e) through the net loss rate *A*(*θ*) which we show for the simulations in Figures 8b-d. We immediately see that *A*(*θ*) has a distinct emerging angular profile, despite cell division and extrusion in the model incorporating no spatial information. In particular, we note that for small values of the alignment strength *J*, where the system cannot form a spiral, there is a strongly positive *A* at large *θ*, near the limbus. This corresponds to a ‘traffic jam’ of cells stuck near their place of birth that are extruded before being able to migrate up the surface. In contrast, for spiral forming configurations, *A <* 0 near the limbus, signalling an enhanced birth rate and lower extrusion rate is balanced by a higher extrusion rate near the centre. Additionally, these *A* profiles are also explicitly dependent on the corneal radius.

**Figure 8.**
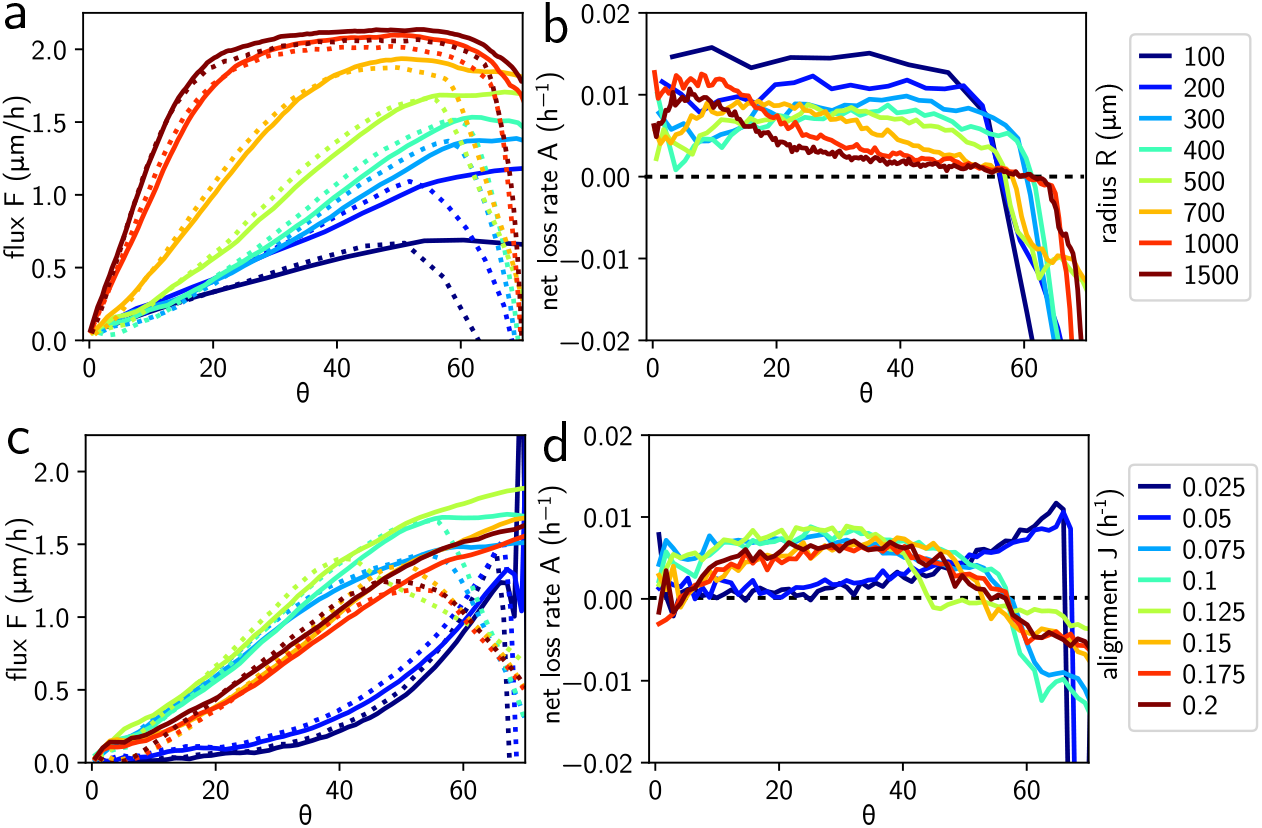
Flux on the simulated corneal surface, and XYZ hypothesis. **a** Radial flux F as a function of polar angle *θ* for different simulated corneal radii *R* at *J* = 0.1 h^−1^. Dashed lines: predicted flux from eqn. (5) using **b** the measured difference between cell division and extrusion rates *A* as a function of *θ*. **c** Radial flux emerges as a function of alignment strength *J* at *R* = 500 µm, and is consistent with (dashed lines) the prediction from net loss rate *A* in **d**, where the elevated loss rate near the limbus for the non-aligning system is clearly visible.

We write an equation for the inward flux *F* (*θ*) = *υ*(*θ*) cos(*α*(*θ*)), i.e. the radially inward component of the velocity. Integrating over the polar angle *θ* from the centre outwards then gives a relationship between the flux at angle *θ* and the integrated loss rate inwards of this point,

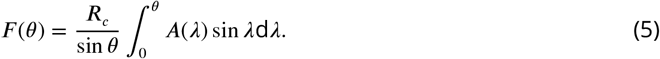

In Figures 8a and 8c, we show the predicted *F* (*θ*)/*R* (dashed lines) compared to the empirically computed flux in the simulations, showing a very good fit. We can clearly see that the flux is also a nontrivial function of both *θ* and cornea radius *R*, explaining the change in spiral shape that we observed as a function of angle *θ* and radius *R*. This dependence on *R* is both explicit in eqn. 5 and due to the nontrivial form of *A*(*θ*), a quantity that we cannot currently measure, but that may at some point become accessible experimentally. We do nevertheless find several interesting constraints on spiral shape: As long as *υ*(*θ*) is nonzero at the corneal centre (and there is no reason for cells to stop moving there), our model implies that *α*(0) = ±*π*/2, i.e. that the spiral centre is a pure vortex. At the same time, since there is nonzero flux inward from the limbus, |*α*(*θ*_*L*_) | *< π*/2, i.e. there has to be an inward component to the spiral at the corneal edge. The *α* = 0 we observe in experiment and simulation for large *R* is simply the most extreme realisation of this.

Since we have established that our model is a polar active material, it should in principle be possible to derive continuum hydrodynamic equations for it, in a similar approach taken for in vitro systems (***Guillamat et al., 2022***). The delicate density-dependent balance that links division, death and velocity magnitude in our system however complicates this: the appropriate symmetry class are the ‘Malthusian’ flocks (***Toner, 2012***), which include division and death. However, the strong approximations inherent in this approach, most notably that *A* = 0 and that *υ* is constant, independent of cell density, make them insufficiently detailed to explain the corneal flux. A future fully quantitative hydrodynamic description of the cornea will require a more sophisticated approach.

### Robustness of the spiral migration pattern and requirement for limbal stem cells

We modelled the cornea in the absence of limbal epithelial stem cell proliferation (Figure 9), first keeping all other parameters as in Figure 6 (left column). The cornea failed to make radial striping patterns. The epithelial pre-pattern of a patchwork of blue and white was never replaced (top row), and multiple +1 and −1 defects persisted long-term (middle row). With increasing J, more pronounced swirling patterns of locally aligned cells were predicted and indeed observed, but without the consistent global patterning of the central spiral - compare the defect tracks in Figure 9g to those of the original system in Figure 6e. We now observed velocity fields that were mostly parallel to the limbus edge, i.e. *α*(*θ*_*L*_) = ±*π*/2, consistent with the absence of any flux into or out of the cornea. Topologically, the states that we do observe, with multiple +1 defects, are possible because we now have singularities at the limbal boundary, consisting of places where the flow field flips from running clockwise to counterclockwise along the edge. This is similar to the multi-defect states with complex boundary behaviour that have been observed for polar active filaments in polygonal confinement (***Kurjahn et al., 2024***), but surprisingly different from the in-vitro systems on a disk - especially the similarly large and polar systems of (***Lång et al., 2024***). We speculate that e.g. higher cell motility and less cell division / extrusion in these systems lead to a different balance between polar motion and effective tissue compressibility. For us, it appears that it is pronounced local cell division and resulting outward flux from the singularities that impedes their merger into the defects and thus disappearance. Note the white patch consisting of a single clone visible in Figure 9f descended from cells at the boundary singularity visible in Figure 9e. We conclude that in vivo, the high levels of cell proliferation in the inner limbus are essential for generating spiral cell motion and so effective corneal resurfacing.

**Figure 9.**
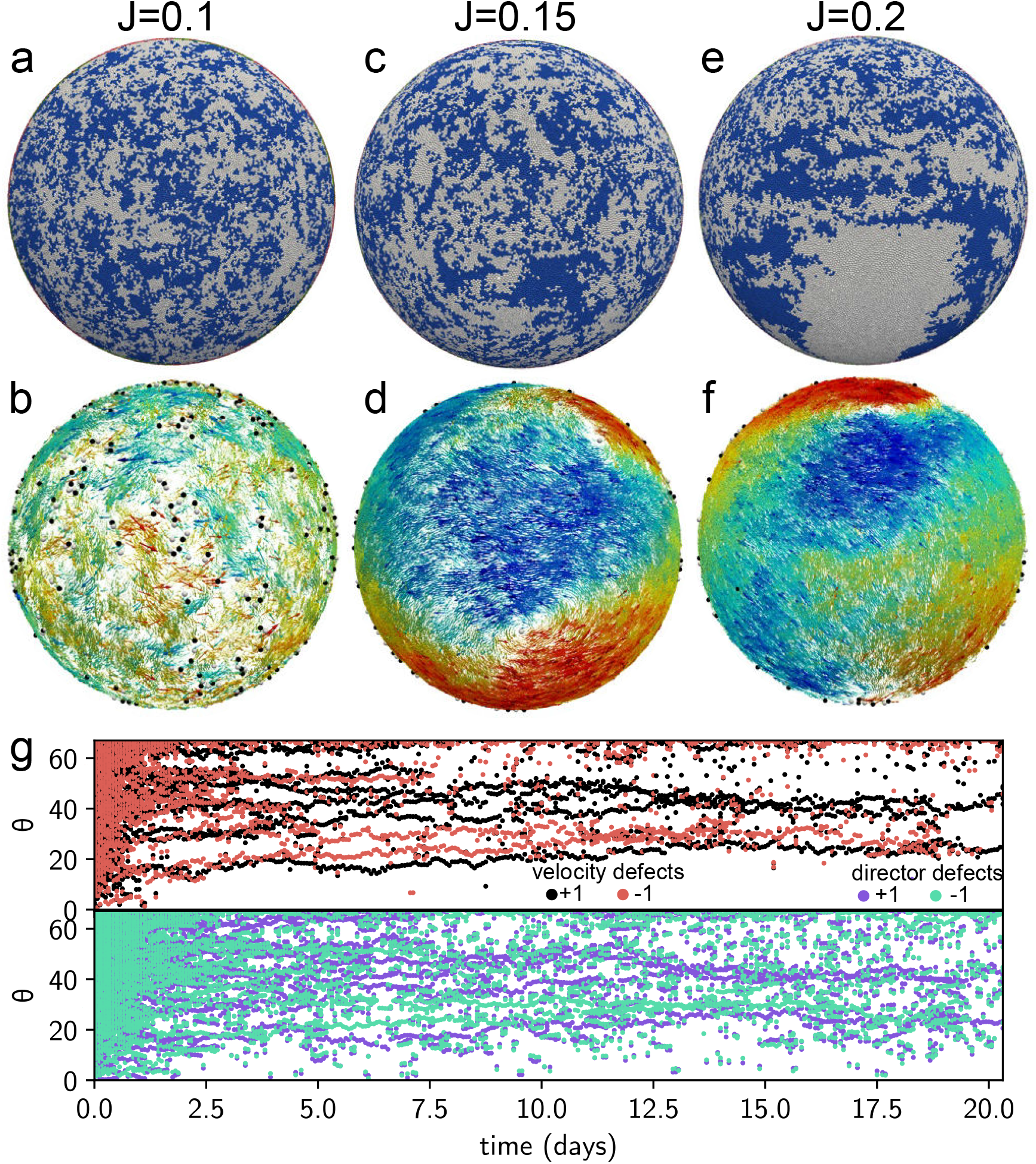
Importance of the limbal stem cells for spiral formation. Result of simulations without limbal stem cells, at *R* = 700 and different *J*, with the TA cells on top and the hydrodynamic flow fields **v**^H^ of the cells at the bottom. **a-b** *J* = 0.1, the direct equivalent of Fig. 6, **c-d** *J* = 0.15 and e-f *J* = 0.2. **g** Defect traces for velocity (top) and director field (bottom) for *J* = 0.15. None of these systems forms a central spiral.

Corneal shape changes significantly between animal species, and thus it is interesting to understand the behaviour of our model in other conditions. We repeated simulations for corneas of different geometries, including a flatter 30 degree cap angle (mimicking human), an oblate ellipsoid (mimicking some ungulates such as sheep), a prolate ellipsoid (mimicking human keratoconus pathology) and also geometries that do not occur in nature - a completely flat cornea (which however resembles in-vitro systems) and a bulging 120 degree cap angle cornea (Figure 10). Radial striping and a single spiral eventually established themselves on all of the modified surfaces. Only in the ‘sheep-like’ oblate ellipsoid there was a tendency for the spiral defect to remain close to the corneal periphery, but there are no data showing whether this reflects an in vivo situation. In Figure 10d, we show defect tracks for the prolate ellipsoid system, which showed the central spiral establishing itself quickly in a very similar manner to the mouse geometry in Figure 6e. While the geometry of these systems varies, topologically they are all the same, with Euler characteristic *χ* = 1 and a boundary condition where there is a net influx of cells from the limbus. Therefore, while the theory for the flux would need to be adapted to the distorted geometry, the general conclusions that at the limbus |*α*(*θ*_*L*_) | *< π*/2 and that the spiral is a pure vortex at its centre remain valid. We conclude that, independent of precise geometry, the combination of topology and cell influx at the limbus robustly leads to spiral formation and effective corneal resurfacing.

**Figure 10.**
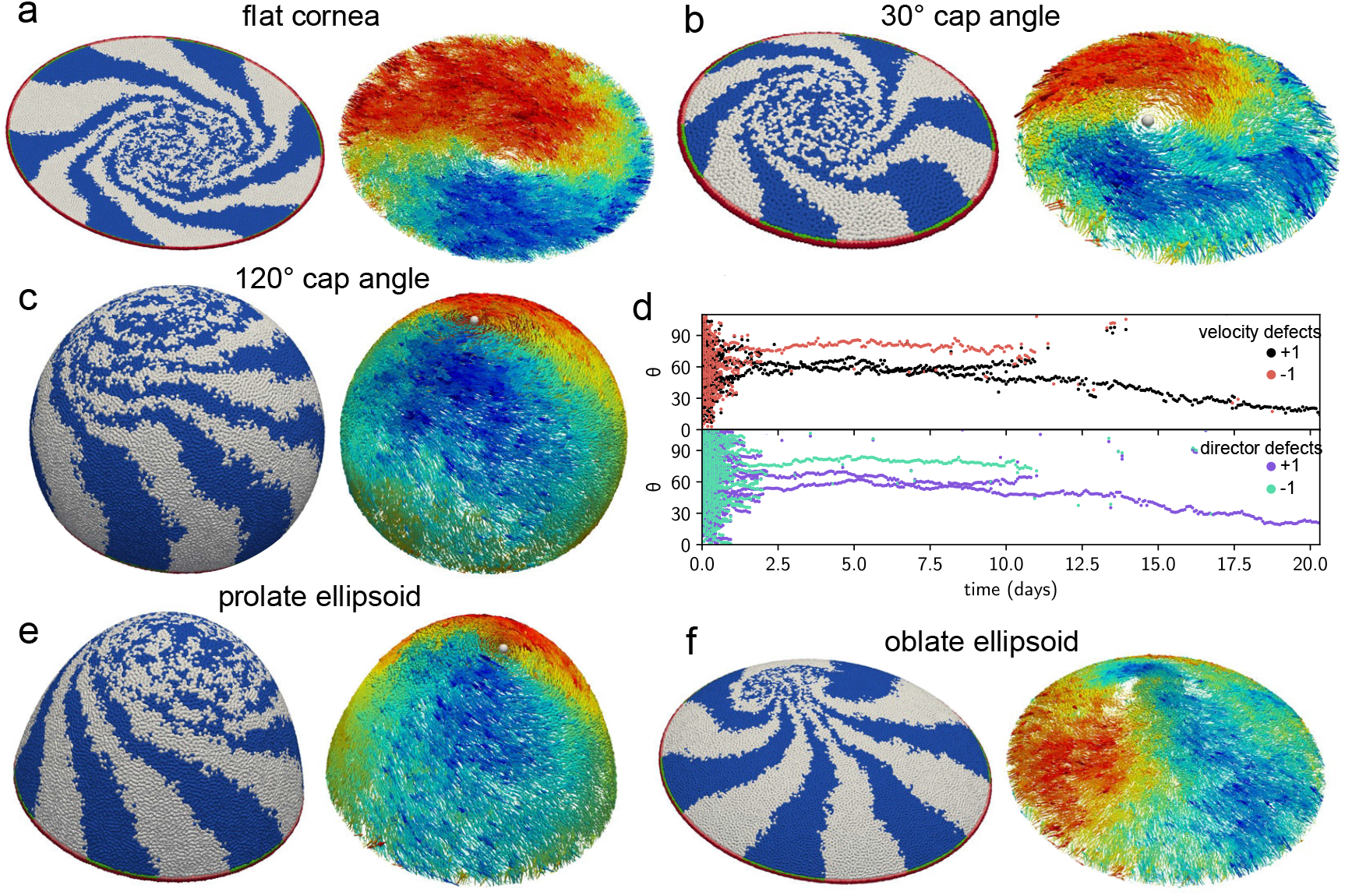
Migration pattern in other geometries, with the ‘blue’ and ‘white’ TA cells marked on the left, and on the right hydrodynamic flow fields **v**^H^ of the cells, representation and other parameters as in the main model and Figure 6. **a** Flat cornea, *R* = 700 µm. **b** Spherical cornea with a cap angle of 30°, *R* = 500 µm. **c** Spherical cornea with a cap angle of 120°, *R* = 300 µm. **d** Defect traces for velocity (top) and director field (bottom) for the system in panel e. **e** Prolate ellipsoidal cornea (radii *a* = *b* = 300 µm and *c* = 500 µm). **f** Oblate ellipsoidal cornea (radii *a* = *b* = 500 µm and *c* = 300 µm). We observe stripe and spiral formation in all systems.

**Figure 11.**
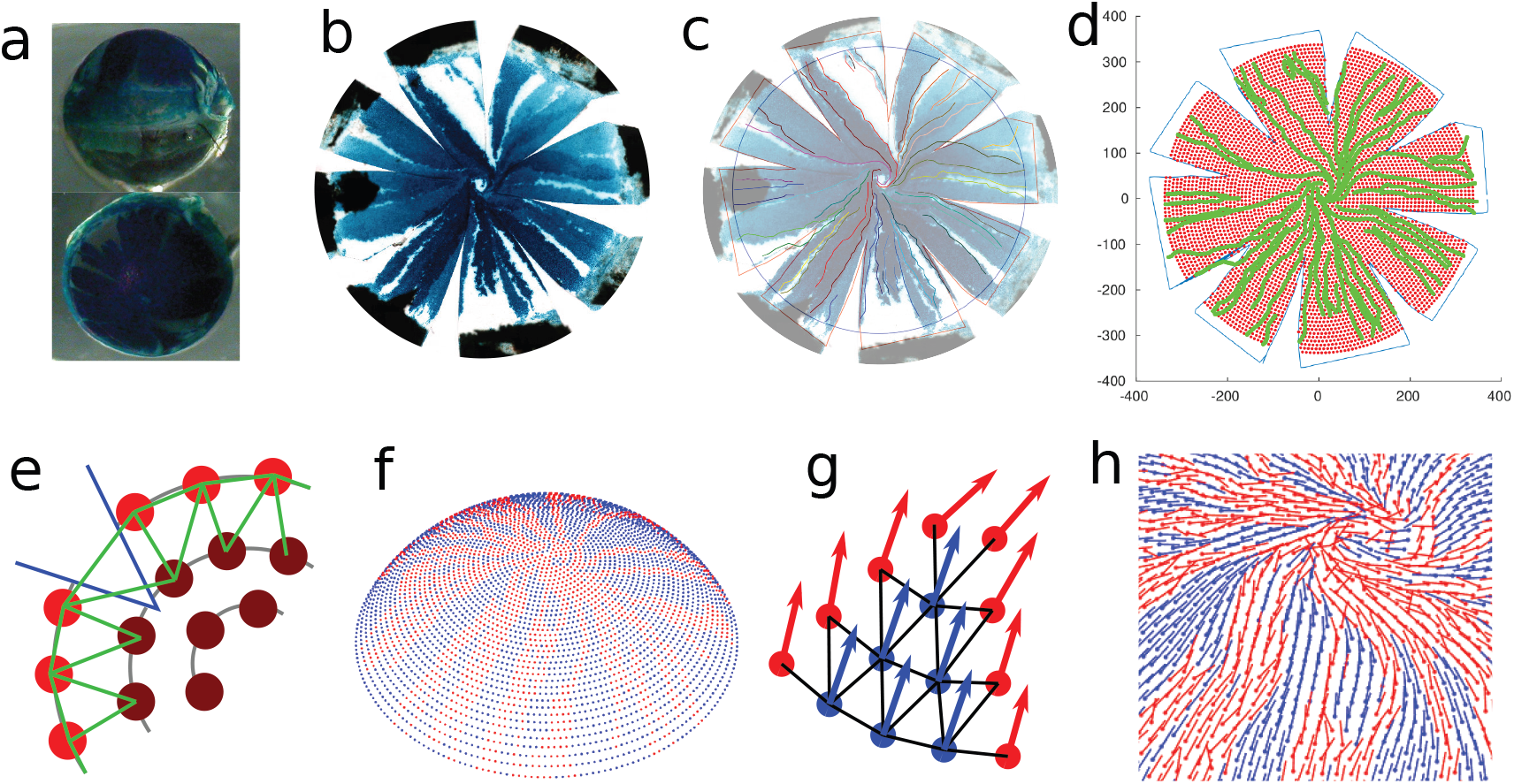
Extracting digitised flow fields from fixed eyes. **a** Side (top) and top (bottom) view of a fixed mouse eye, stained for LacZ. The line of the limbus is clearly visible. **b** Dissected, flattened cornea, note radial cuts and the clearly visible stripes. **c** Using Inkscape, we used different colours to draw (1) the cut boundaries (2) a circle at the position of the limbus and most importantly (3) all visible stripe edges as new layers on the image from (**b**). **d** After processing this drawing with MATLAB, we obtained (1) the outline of the dissected tissue (2) the location of the cornea and (3) the stripe locations. We placed 40 concentric rings of evenly spaced points on the cornea, omitting the cuts. **e** ‘Moulding’ the cornea: Starting from the centre and moving outwards, we relaxed each ring by equilibrating the springs indicated in green; a cut location is outlined in blue. **f** Location of the points defined in (**d**) on the cornea after ‘moulding’. **g** Using the local tangent to the stripe edges, we inferred the velocity direction on points coloured red (red arrows). We then relaxed the director at points without stripe edge (blue) assuming director alignment between neighbours (black lines). **h** Final estimate for the flow direction of the cornea, including central spiral defect.

**Figure 12.**
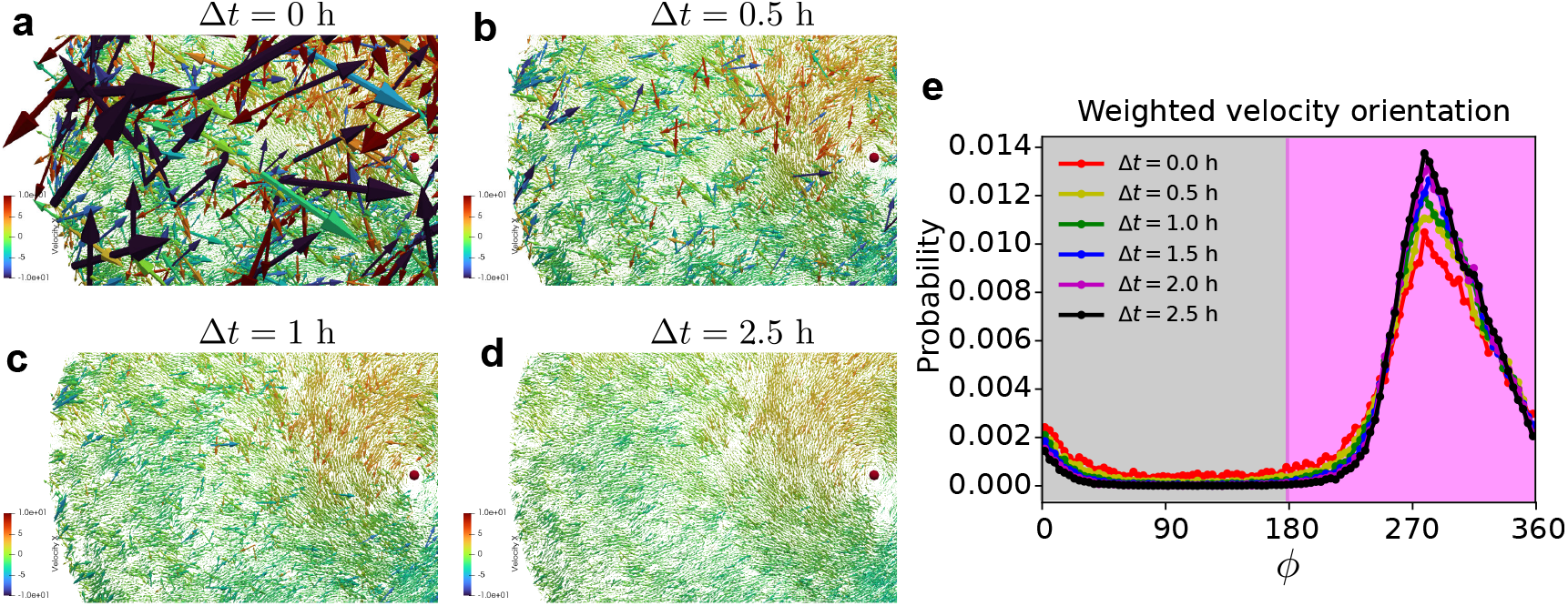
Emergence of the hydrodynamic velocity. **a** Instantaneous velocity arrows on the *R* = 700*µm* cornea shown in Figure 6, colored by x (left/right) component of the velocity. **b-d** Hydrodynamic velocity arrows defined as 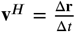for increasing time intervals Δ*t*, showing convergence to smooth field after Δ*t* = 2.5*h*. **e** Distribution of the angle *ϕ* of the velocity vectors (as defined in Figure 3), weighted by velocity magnitude, showing increasingly sharply peaked inward (pink) motion as Δ*t* increases.

## Discussion

### Conclusion

We introduced an in silico agent-based model that describes the complex collective motion pattern observed in the corneal epithelium as an emergent phenomenon. The model only requires simple assumptions about local cell behaviours and neither relies on any organ-scale pre-patterning or regulation nor requires the imposition of any external guidance cues or any shear and tensile strains that may otherwise cause the sliding of epithelial cells in logarithmic vortices (***Mohammad Nejad et al., 2015***). In vivo, cell migration patterns can, therefore, be quantitatively explained by a simple agent-based model commonly studied in the physics of soft active matter systems. This is the first corneal model to successfully reproduce the central vortex motion of epithelial cells around the corneal centre in simulations. The key outcomes from this study are that to recapitulate experimentally observed patterns of cell migration in vivo, cells need to: 1) be able to actively migrate, 2) respond to mechanical pressure due to the population of other cells, 3) align their migration direction to that of their neighbours (i.e. ‘go with the flow’), 4) have density-dependent control of proliferation, and 5) maintain a high rate of limbal cell proliferation. Then the emerging pattern is robust to noise, i.e. stochastic components in cell motion and proliferation, and corneal size and shape. Furthermore, we were able to determine model parameters from experiments, which allowed for quantitative comparison between the in silico model and the biological system. Changing model parameters results in markedly different cell migration patterns that may apply to other biological scenarios, underscoring the power of the agent-based active matter models in describing collective cell behaviours.

Notably, in the cornea, where motile cells are bounded by immobile stem cells, the model predicts that at no time do cells need guidance cues to direct them towards the corneal centre. Given the step into the cornea from the limbus, which we impose on the simulations and reflects the likely durotactic pressure on cells moving from the soft limbal stroma to the stiffer corneal stroma, the radial centripetal migration that has long fascinated biologists can then be explained as being generated by a biophysical model.

The high parameter of limbal cell division that is required to produce robust radial striping is superficially at variance with the paradigm that suggests limbal epithelial stem cells to be slow-cycling. However, within the inner limbus, cycling stem cells give rise to rapidly dividing TA cells. Although up to 25% of limbal cells may be slow cycling, we previously calculated that the mean limbal epithelial cell cycle time is lower than that observed in the corneal epithelium (***Sagga et al., 2018***). There is, therefore, no discrepancy between the stem cell parameters imposed on our model and the real biological situation. We also tested the consequences of the removal of limbal epithelial stem cells from the simulated system and showed that in these circumstances the pinwheel pattern of cell migration does not occur and the cornea maintains multiple unresolved +1 and −1 discontinuities (Figure 9). This is biologically relevant as it has been shown that at least some of the corneal epithelial TA cells have high clonogenic potential and in some species, the corneal epithelium can be maintained for a substantial period of time after limbal epithelial stem cell ablation. Our simulations suggest that the corneal epithelium indeed should not be expected to immediately degrade upon loss of stem cells. However, the pattern of cell migration across the cornea in those circumstances would be highly irregular. Such a cornea may not be able to repair wounds effectively.

Analysis of the agent-based simulations data shows that at the start of the simulations, the sheet is peppered with ±1 topological defects, but those collide with each other and cancel out until a single +1 defect persists. Across several simulations, the model also found multiple +1 and −1 defects during the first 5 days, but by about simulated day 10 invariably only one defect remained. During simulations, each topological defect moved stochastically, which enabled all but one +1 defect to collide and eventually cancel out. This ‘Highlander’ model, where only one defect can survive, is a direct result of simulations but also has strong relevance to our understanding of biology. The start of the simulations represents the young mouse corneal epithelium which consists of a random unoriented patchwork of ‘blue’ and ‘white’ cells, full of topological defects, with the development of the single central vortical swirl only manifesting after days or weeks of radial cell migration (***Collinson et al., 2002***). The simulation suggested that it may be possible that two or more +1 discontinuities survive long-term (providing the overall topological charge of the cornea remains +1) if one or more −1 and +1 discontinuities happen not to collide. An in vivo example of this is seen in Figure 1. We have observed persistent recurrence of +1 and −1 discontinuities in disease states, such as Pax6-heterozygous mice in which the corneal epithelium is fragile and subject to persistent abrasion (***Collinson et al., 2004***). Discontinuites are also created after peripheral wounding of the cornea, and these may persist long term if widely separated (Sagga et al., in prep).

### Outlook: Models of the cornea and active matter

The physics of active matter has emerged as a powerful tool to describe many collective processes in biological systems, revealing that many complex biological processes rely on common physical mechanisms (***Alert and Trepat, 2020***). The cornea studied here is one such example. What, however, sets it apart from most other active matter studies of collective cell migration is the presence of curvature. Curvature alters the common notions of straight and parallel lines, which affects how active agents interact with each other if confined to move on a curved surface. It can also lead to the appearance of topological defects (e.g. the +1 defect at the cornea top), which can have important implications for the development of the organism, e.g. as was recently argued to be the case for hydra (***Maroudas-Sacks et al., 2021***). Compared to the efforts put into understanding active behaviour in flat spaces, the current understanding of the interplay of curvature and activity is, however, very limited. We, therefore, hope that this work will motivate further research in this direction.

We also remark that the mutual annihilation of integer topological defects as the system ages has recently been reported in an in-vitro system immortalized human keratinocytes confined to a disk and confirmed in agent-based simulations (***Lång et al., 2024***). The model proposed there in fact uses the same combination of overdamped dynamics and self-alignment as ours, and is thus in the same class of active polar matter. In contrast, however, it is an active *solid* model without cell division and extrusion, and indeed, any cell rearrangement at all, and model results can therefore not directly be compared to ours.

Previously, coordinate-based in silico models of ocular striping have attempted to model the cornea, but did not adequately recapitulate the biological state, due to not accounting for the correct geometry and curvature of the cornea (70° hemisphere) and often ignoring factors such as cell migration. In one study, the cornea was modelled on a flat surface with population pressure from neighbouring cells being solely responsible for mobility, with cells moving from higher to lower pressure (***Lobo et al., 2016***). In contrast, another study used a simple stochastic approach with a hemisphere geometry, in which patterning is driven by proliferation, ignoring migration (***Moraki et al., 2019***). Both studies explored the effects of manipulating the amount of cell loss in different regions from the surface of the cornea through higher terminal differentiation of the TA cells (e.g. mimicking UVR damage), employed lineage tracking and the ability of all the cell types to undergo asymmetric or symmetric divisions. Neither model incorporated all the components of cell-cell interaction, persistence, correlation and noise, and neither was able to generate a spiral migration pattern, a key feature observed in vivo.

### Outlook: The cornea and cell migration

An important feature of our model is its stability over scale and corneal shape, reflecting the fact that patterns of radial migration have been observed in multiple species, including humans, with very different corneal shapes and sizes. It also allows us to directly test hypotheses for corneal homeostasis and make predictions about the corneal response to genetic, pharmacological and physical manipulation.

Eye opening in mice, at around 10 days postnatally, predates the emergence of radial striping patterns in vivo. The increase in corneal abrasion and cell loss concomitant with eye opening is presumably the trigger for limbal stem cell activation, though this has never been experimentally demonstrated, and the radial orientation of cell movement and neuronal projection is observed relatively rapidly following eye opening (***Iannaccone et al., 2012***). We suggest that the increase in cell loss associated with eye opening is a wound mimic which drives the rapid initiation of radial stripe patterns in young mice, and this is supported by the in silico modelling. By changing the parameters of simulated cell behaviour, it will be possible to mimic the effect of mutations in genes controlling cell velocity (e.g. Pax6) and polar alignment (e.g. Vangl2), both of which have been shown to disrupt patterns of corneal epithelial migration in vivo.

One aspect of the corneal system that was largely ignored in this study is the relationship between corneal epithelial cell migration and the radial projection of sensory axons between those cells (Figure 1b, c). The most intuitively parsimonious explanation of corneal anatomy would suggest that the axons tend to follow the path of least resistance within the epithelial flow. However, experimental observation shows that radial axon projections appear in the mouse corneal epithelium from about 3 weeks of age (***McKenna and Lwigale, 2011***), i.e. before epithelial striping becomes apparent in transgenic LacZ and GFP reporter mosaics at about 5 weeks (e.g. (***Collinson et al., 2002***; ***Iannaccone et al., 2012***). This suggests the possibility that the corneal epithelial cells may be guided by a pre-existing alignment of the sensory axons. However, the appearance of radial projections of neurons before the development of epithelial stripes is easily explained because experimental observations of stripe formation show (***Collinson et al., 2002***), and our model predicts (Figure 6a, b) that epithelial cell migration proceeds for many days before the resultant striping patterns become apparent in transgenic reporter assay mice. The observed appearance of radial axon projections before the lagging epithelial striping patterns is fully consistent with the biological scenario that the axons follow the migrating epithelial cells.

This does not of course preclude the possibility that guidance cues, e.g. contact with axons, exist that modulate or refine the pattern. (***Walczysko et al., 2016***), show that isolated epithelial cells in tissue culture still migrate with a slight but significant radial bias when plated onto the deepithelialised (and de-nervated) corneal stroma. Clearly corneal epithelial cells respond to their physical environment (as evidenced also by the component of alignment in our model). Since the radial striping at cornea centre in most of our simulations was not as tidy as in real life, it is very possible that some form of guidance cue refines the pattern.

Understanding corneal epithelial cell homeostasis is essential to improve wound healing after corrective eye surgery, and to develop treatments for corneal diseases. The cornea is a paradigm system of epithelial maintenance by a population of stem cells. It is a system of choice for interdisciplinary studies of the biomechanics of cell migration because it is simple (few cell types), transparent (thus accessible to in vivo imaging), nonessential for life (thus amenable for mutation), and accessible to modelling. However, the principles and mechanisms of the cell migratory response are likely to be similar to those that underlie cell migration patterns in other tissues that are morphologically or functionally less amenable to investigation. The findings here, that a dense active matter model can recapitulate and predict patterns of epithelial cell behaviour in vivo, likely have general relevance to multiple cell migration events throughout embryogenesis and wound healing. The morphogenetic movements and epithelial-mesenchymal interactions that produce swirls of hair follicles on the head or that underlie patterns of pigmentation commonly associated with ‘Blaschko’s lines’, may also be predicted by an active matter model (***Plikus and Chuong, 2004***; ***Kücken and Newell, 2005***; ***Tenea, 2017***).

## Materials and Methods

### Mice

We obtained all animal tissue samples from the Medical Research Facility of the University of Aberdeen in accordance with the Animal (Scientific Procedures) Act 1986. H253 mice which carry an X-linked LacZ transgene, subject to X-inactivation (***Tan et al., 1993***), were maintained on a CBA/Ca genetic background, breeding transgenic males with wild-type CBA/Ca females to derive hemizygous, LacZ-mosaic females. We stained eyes from adult female mice at least 15 weeks old for the flow pattern and cell migration work.

### Whole corneal imaging

We collected, fixed and stained LacZ-mosaic eyes with X-Gal (***Collinson et al., 2002***). We dissected the corneas from the eyes and added eight radial cuts from the edge to near-center such that they could be flat-mounted on a microscope slide with glycerol and a coverslip applied. Following flat-mounting the corneas were imaged using a slide scanner.

### Immunohistochemistry

We used mouse corneas for immunohistochemistry to visualise cells undergoing mitosis and determine the location of the centrosome. Mouse eyes were fixed in 4% paraformaldehyde for 2 hours, washed in phosphate-buffered saline (PBS), then we dissected the corneas and post-fixed in methanol for 1 hour at −20 °*C*. After 3, 20 minute washes in PBS, corneas were immersed in blocking buffer (PBS, 4% normal donkey serum, 0.3% bovine serum albumen, 0.1% Triton X-100) for 1 hour at room temperature. Primary antibody [rabbit anti-phosphohistone-H3 (06-570, Millipore) or rabbit anti-pericentrin (ab448, abcam)] was dissolved 1:200 in blocking buffer and added to corneas overnight. After 3, 20 minute washes in PBS, secondary antibody [Alexa-488-conjugated donkey anti-rabbit (Molecular probes)] was dissolved 1:200 in blocking buffer and added to corneas for 2 hours at room temperature in the dark. After 3, 20 minute washes in PBS, corneas were flat-mounted as above in a drop of Vectashield with DAPI (H-1200, Vector Labs) for fluorescence microscopy.

### Estimating velocity fields from stripes

The mouse cornea is not a full hemisphere so we measured the spherical cap angle and eye diameter experimentally in nine adult corneas giving an average of 71.1° ± 0.3° SEM cap angle, and 3.60 ± 0.02 mm SEM in diameter, rounded to 70° and 3.6 mm, respectively for modelling.

In vivo, we inferred three-dimensional corneal epithelial flow patterns by the proxy of tracing the boundaries between LacZ-positive and negative radial stripes in XLacZ mosaic corneas. As shown in the in silico model, despite some small-scale mixing at the boundaries, the steady-state flow field is parallel to the stripe boundaries. We proceeded as follows to extract approximate flow patterns from XLacZ corneal epithelia: first, we captured scaled images of each eye (Figure 11a). After photography, we dissected the cornea from each eye, made 8 radial partial cuts then flat-mounted the cornea for high-resolution images (Figure 11b). Using Inkscape image processing software (***Inkscape Project, 2020***), we drew a circle corresponding to the limbus on the image in a first colour, together with the edges of the radial cuts in a second colour, generating a background image file. In a second layer, we drew the edges of the stripes. Because different clonal stripes were often in different shades of blue, it was usually also possible to resolve clonal borders within blue stripes, with as much detail and covering as possible (Figure 11c). Different colours were used to distinguish separate flow lines, and we saved this layer as a separate image file.

These digitised corneas were used to extract the flow fields using a custom MATLAB code: We first read the background image file and extracted the limbal circle as well as the edges of the cuts. We then placed equally spaced points in 40 concentric rings in only the interior of the irregular polygon of the actual cornea defined by this method (Figure 11d). On each ring of radius *r*, we defined *n* individual points with spacing *dr* = 2*πr*/*n* with coordinates **r**_*k*_ = (*r* cos *α*_*k*_, *r* sin *α*_*k*_), where *α*_*k*_ = 2*πk*/*n*; here the *k* corresponding to coordinates inside the cut areas were removed. The remaining points were where we then aimed to define the flow field.

We then read the image file containing flow lines and extracted the pixels corresponding to drawn lines, mapping each colour to a different label. We fitted directional unit vectors to pieces of lines using linear regression on groups of pixels from the same line. Here we chose the direction of the arrow so that arrows at the limbus pointed inward, and so that subsequent arrows on the same line were in the same direction. We applied an error correction step so that velocity vectors of adjacent ring points were required to point in the same direction, flipping individual directors if necessary. Finally, we averaged orientations over the different arrows within 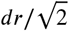 distance of a point to define the velocity unit vector on ring points. These inferred velocity vectors are shown as red arrows in Figure 11.

Since the cornea corresponds to a spherical cap with an approximate 70° angle, we needed to close the gaps left by the radial cuts. To achieve this, we applied a virtual moulding technique to the image points starting from the inside and moving down ring by ring (Figure 11e): To move the tissue on either side of the cuts together, we introduced springs between all neighbouring pairs of points, and also between points between the ring and the previous (interior) one if they were within distance *dr*. We defined an energy function for our network of springs,

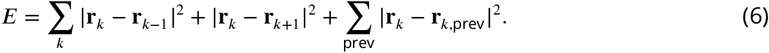

Using a Monte Carlo simulation with effective temperature *kT* = 0.01, we then sequentially equilibrated each ring for 1000 Monte Carlo sweeps, starting from the centre. The final angles were then equally spaced, closing the cuts. The final arrangements of points on the now curved cornea are shown in Figure 11f. We then inferred the flow direction in the region between stripe edges: Since neighbouring flow fields (or ‘spins’) are expected to be similar to each other, we introduce an aligning energy term akin to modelling magnets, i.e. the XY model, 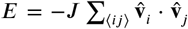, where the sum is over neighbouring points only (black lines in Figure 11g). In spherical coordinates, (ê_*r*_, ê_*θ*_, ê_*ϕ*_) as shown in Figure 2c, we could define the orientation of the flow direction 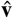 through the deviation angle *α* from inward flow, i.e. 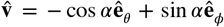 in the local tangent plane to the sphere. One could then show using parallel transport (to first order), that the dot product corresponds to 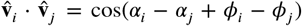. Here *α*_*j*_ was shifted by the difference in longitude *ϕ*_*j*_ − *ϕ*_*i*_ between the two points, which is significant for direct neighbours close to the pole. We then again used a Monte Carlo approach with simulated annealing where we gradually reduced the effective temperature in steps from *kT* = 1 to *kT* = 0.001 to minimise the energy of our spin configuration while keeping the red arrows where we already knew the flow direction was fixed. As the directions are parameterised by declination *α*_*i*_, all arrows remain tangent to the sphere, ensuring that our flow field is on the corneal surface, not into it or out from it. The resulting estimated flow field is shown in Figure 11h, and in main Figure 2a.

### Defect finding algorithm and spherical coordinates

In order to identify the locations of topological defects in the flow field, we first constructed a Delaunay triangulation. For the experiment (Figure 2d), we first projected all points into the xy plane, while for the simulation, we derived the Delaunay triangulation from the convex hull of the particle positions on the sphere. The unit-length vectors along the direction of velocity can be written using the angle *θ* with an axis in the local tangent plane, i.e. 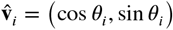. We located topological defects by summing over the angle differences around the loop of the projection plane if one traverses each triangle counterclockwise. The topological charge associated with each triangle is 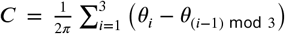 (***Kamien, 2002***; ***Henkes et al., 2018***). This is an approximation of the Burger’s loop integral 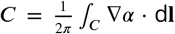 d**l**, and allowed us to locate defects as the centres of loops with *C* ≠ 0. Generally, *C* = 0 (e.g. in loop A) for all triangles except the one at the centre of the spiral, where *C* = 1, corresponds to the central +1 topological defect (loop B). To obtain the spiral angle profile *α*(*θ*), we first located the topological defect(s) in the 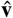 field. As shown in Figure 2a (experiment) and Figures 6c and 6d, we reliably found a single +1 topological defect near the centre for mature corneas. We then shifted our spherical coordinate system so that the defect was at the pole, and defined the corneal spiral angle *α*(*θ*) through averaging *α*= arctan 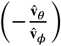 over bins of latitude *θ*.

### Time-lapse imaging

We dissected corneas, halved them, sterilised them in 1:10 penicillin/streptomycin solution for 10 minutes then washed them in PBS before attaching the halved sections, stromal side down, to uncoated 60 mm tissue culture dishes and flooding them with corneal culture medium following (***Hazlett et al., 1996***). Explant cultures were maintained at 37 °*C*, 5% CO_2_ until imaging. Cultures were imaged in HEPES-buffered media using phase-contrast optics on an inverted Leica (DM IRB) chamber microscope for 24 - 48 hours, taking images at 10-minute intervals for PIV analysis.

### Division and centrosome angles

We used ImageJ to measure mitotic angles of division (phospho-histone-h3, DAPI) and centrosome position (pericentrin, DAPI) in relation to the centre of the eye (Figure 3).

### Computational simulation details

Individual corneal epithelial TA cells obey the following equations of motion,

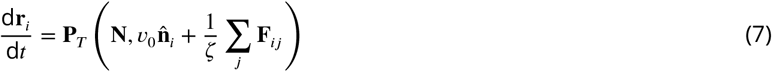

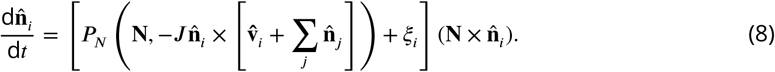

The first equation relates the time derivative of the position, **r**_*i*_, of the i^th^ cell to the active self-propulsion velocity *υ*_0_ along a migration direction described by the unit-length vector 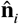. **F**_*ij*_ is the elastic force with the neighbouring cells labelled by indices *j*, and forces are summed over all neighbours within the interaction range. The constant *ζ* is the friction coefficient between cells and the substrate (i.e. the stroma). The second equation describes the rotational dynamics of the vector 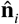. In addition to the term *ξ*_*i*_ that accounts for random changes in the direction of 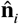, i.e. the rotational diffusion, there is a term that describes alignment with directions of neighbouring particles 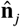, and a term that models alignment of the particle’s direction with the direction of its velocity 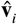. The rotational diffusion is assumed to be the simple white noise with zero mean and variance 2*D*_*r*_. Finally, the projection operator **P**_*T*_ (**N, a**) = **a**−(**N**⋅**a**)**N** enforces that the motion remains in the plane orthogonal to the local normal 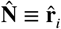, the projection operator 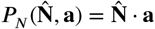 projects the rotation axis of 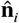 to be along the local normal vector. In isolation, on the sphere, this model has a transition to a flocking ring state at sufficiently large *J*. A full analysis of this model is given in (***Sknepnek and Henkes, 2015***).

For our simulations to match corneal ‘explants’ and tissue culture ‘plastic’ colonies, we used the same model on a flat plane, so that 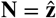, simply the out-of-plane direction. If we define the polar angle *ϕ*_*i*_ through 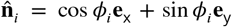 and the velocity angle through **v**_*i*_ = *υ*_*i*_ (cos *θ*_*i*_**e**_x_ + sin *θ*_*i*_**e**_y_), where **e**_x_ and **e** are the unit-length vectors, respectively, along the *x* and *y* axes of the simulation box, the model eqns. (7) and (8) reduce to a simpler form

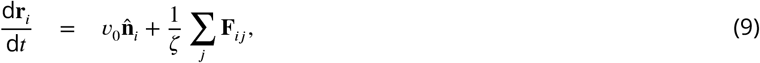

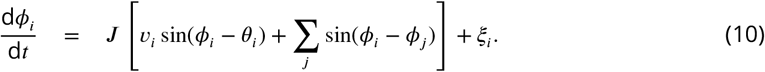

We used a simple, short-range interaction potential that includes both repulsive and attractive parts, together with regulated cell division and removal. This model was introduced and characterised in (***Matoz-Fernandez et al., 2017***). Briefly, the resulting force between particles *i* and *j* with radii *σ*_*i*_ and *σ*_*j*_ and distance vector **r**_*ij*_ = **r**_*j*_ − **r**_*i*_ was given by

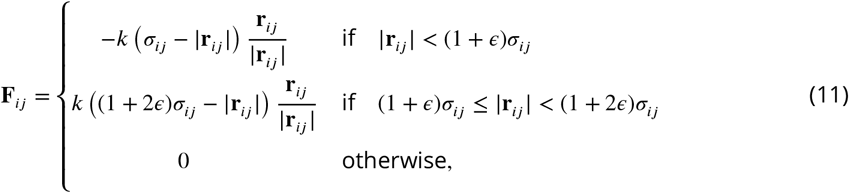

where *σ*_*ij*_ = *σ*_*i*_ + *σ*_*j*_ and the dimensionless number *ϵ* sets the attraction depth and width. Due to the soft potential, this model is unstable for *ϵ* ≥ 0.2, and we chose *ϵ* = 0.15 throughout, as well as 0.3 polydispersity of the cells. Particles (cells) were modelled to divide and disappear from the sheet according to a constant squeeze-out rate *a* but a division rate *d* = *d*_0_(1 − *z*/*z*_max_) that depends on the local density through *z*, the number of neighbours of the cell (defined as those in range for **F**_*ij*_ ≠ 0). As shown in (***Matoz-Fernandez et al., 2017***), there was a ‘self-melting’ dense liquid steady state in this model below a threshold *a*/*d*_0_ ≈ 0.1, i.e. where extrusion is sufficiently rare for cells to divide to reach confluence at *z* ≈ 6 and so in turn inhibit division. The parameters we used for the simulations are summarised in Table 1. All simulations reported here were carried out in our simulation package (***SAMoS, 2024***) using custom configuration and input files. All analysis of simulation data, including coarse-grained fields, topological defects and angular averaging were carried out using our analysis package (***SAMoSA, 2024***).

### Obtaining coarse-grained fields from simulation

Figure 12 shows the emergence of the hydrodynamic velocity field *υ*^*H*^ upon time averaging.

### The XYZ model revisited: details

Consider a small patch of area *S* of corneal tissue containing *N* cells at time *t*. One can write the change of number of cells between times *t* and *t* + Δ*t* as

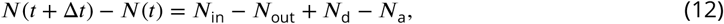

where the terms are given by the number of cells that flow into the patch (*N*_in_), flow out of the patch (*N*_out_), are born (*N*_d_) and are squeezed out (*N*_a_).

This is nothing but a form of a continuity equation which expresses that all cells need to be accounted for, and it can be written in a differential form. If *ρ*(**r**) is the local number density of cells, then *N* (*t*) = ∫_*S*_ *ρ*(**r**, *t*)d**r**. Other terms can be rewritten as functions of *ρ*: The total number of births during time span Δ*t* is given by *N*_*d*_ = Δ*t* ∫_*S*_ *d*(*ρ*)*ρ*(**r**, *t*)d**r**, where *d*(*ρ*) is the density-dependent division rate. Similarly, *N*_a_ = Δ*t* ∫_*S*_ *a*(*ρ*)*ρ*(**r**, *t*)d**r**, where *a*(*ρ*) is the density-dependent extrusion rate. The flux of cells through the border of the patch can be expressed through a number flux current **J** = *ρ***v**, i.e. the number of particles with velocity **v** passing through a unit line in unit time, with units of inverse time and length,

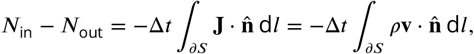

where the integral is over the boundary of *S* and **n** is an outward pointing unit-length normal. Putting all of the pieces together, one obtains

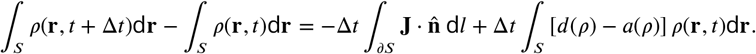

Using the divergence theorem on the first term on the right-hand side, it can be rewritten as 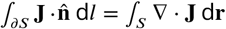, followed by the limit Δ*t* → 0. This leads to the continuity equation

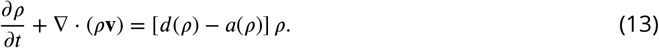

#### Cornea spiral flux equations

We now explore the implications of the continuity equation for an existing spiral flux on the corneal surface. Consider a cornea that is on the surface of a sphere with radius *R*, with an opening angle at the limbus of *θ*_L_ = 70°. Using spherical coordinates as in Figure 2c, with *θ* as the angle from the cornea centre and *ϕ* the azimuthal angle, we assume that we have an established symmetric spiral flux ending at the cornea centre, making an angle *α*(*θ*) with the radially inward direction. We assume that the density is symmetric, i.e. that we have *ρ*(*θ*), and that the average cell velocity in the local polar coordinate system is given by **v** = *υ*(*θ*) [cos *α*(*θ*) ê_*θ*_ + sin *α*(*θ*) ê_*ϕ*_]. We also recall that in spherical coordinates, the divergence of vector **U** = *U*_*r*_ê_*r*_ + *U*_*θ*_ê_*θ*_ + *U*_*ϕ*_ê_*ϕ*_ is given by

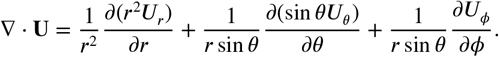

Here, we have **U** = *ρ***v**, and we are always on the corneal surface with *r* = *R*_*c*_ and we eventually write

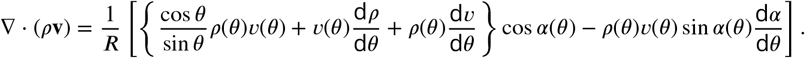

In steady-state, i.e. when the system becomes time-independent so that 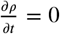, eqn. (13) reduces to ∇ ⋅ (*ρ***v**) − *A*(*ρ*)*ρ* = 0, where the effective net loss rate *A*(*ρ*) = − [*d*(*ρ*) − *a*(*ρ*)]. We then find a time-independent, ordinary differential equation in the variable *θ* linking velocity, density and spiral angle:

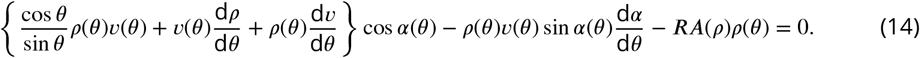

We can make one more simplifying assumption, namely that *ρ* is approximately constant in the system. This is reasonable, first observationally as shown in Figure 13, and second, because the relaxation time of the potential is much much faster than the typical migration time. We will however retain two effects on the net loss rate *A*: First, *A*(*ρ*) is very sensitive to density, so we can not fully neglect its effect. Second, in our system TA cells have a typical lifetime, suppressing their death rate near the emergence from the limbus. We will therefore retain an angle dependence *A* = *A*(*θ*). Then the final continuity (XYZ) equation is given by

**Figure 13.**
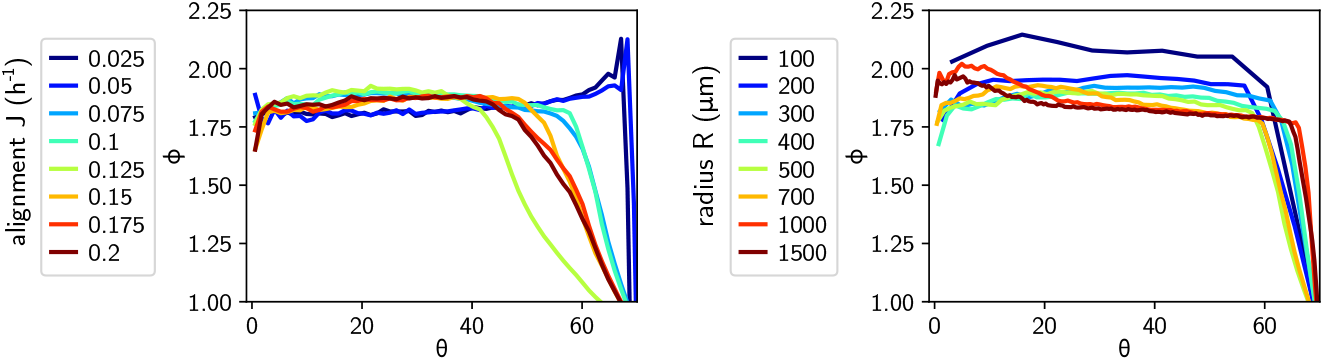
Packing fraction *ϕ* = *πr*^2^*ρ* for the simulations as a function of angle *θ* from the centre, for different alignment strengths *J* and radii *R*. Note that for *J* above the alignment transition and all *R* the apparent regions of lower *ϕ* at *θ >* 50° are due to corneas with asymmetric defect patterns clipping the edge of the cornea at those angles away from the defect.

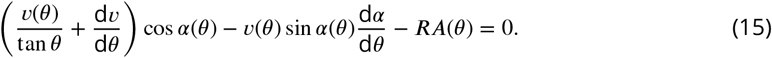

Figures 6 and 7 show the numerically observed spiral angle *α*(*θ*), velocity magnitude *υ*(*θ*) and the effective death rate *A*(*θ*). It becomes immediately clear that none of these functions is simple, and that there is a strong dependence on the radius of the simulated cornea, despite *R* only appearing in the scaling of the net death rate *A* in eqn. (15). A major contribution to the complexity is the form of *A*(*θ*): As can be seen, it is negative near the limbus where cells are recent descendants of stem cells, and then positive and approximately constant towards the inner regions of the cornea. Nevertheless, eqn. (15) is satisfied: the centrally inward (along ê_*θ*_) part of the velocity is given by *F* (*θ*) = *υ*(*θ*) cos(*α*(*θ*)), representing the inward flux of material. Undoing a number of the spherical coordinate derivatives, we can rewrite equation 15 as a function of the inward flux:

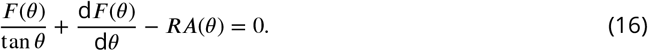

We supplement this equation with the following boundary conditions. First of all, the flux needs to vanish at the central defect, i.e. at *θ* = 0, *F* (0) = 0. This means that either *υ*(0) = 0, i.e. the cells completely stop moving anywhere on average at the centre, or cos(*α*(0)) = 0, which means that the velocity is at an angle *α*(0) = ±90°, or completely orthogonal to the inward direction. We can then in fact find an integral solution for *F* (*θ*): First make the change of variables *u*(*θ*) = *F* (*θ*) sin *θ*, which when replaced in eqn. (16) leads us to

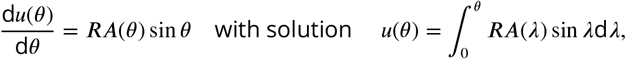

using the boundary condition *u*(0) = *F* (0) sin(0) = 0. Then we arrive at our final prediction for the flux on the cornea for a given death profile *A*(*θ*):

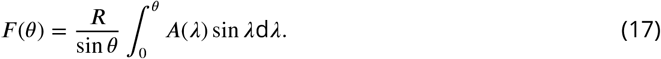

## Supporting information

Description of supplementary videos

## Author details

### Kaja Kostanjevec

- University of Aberdeen, School of Medicine, Medical Sciences and Nutrition, Aberdeen AB25 2ZD, United Kingdom

Contributions: Experimental work. Experimental data analysis. Performed numerical simulations. Wrote and edited the manuscript.

Competing interests: No competing interests declared

### Rastko Sknepnek

- School of Life Sciences, University of Dundee, Dundee, DD1 5EH, United Kingdom
- School of Science and Engineering, University of Dundee, Dundee, DD1 4HN, United Kingdom

Contributions: Model and code development. Edited and commented on the manuscript. Competing interests: No competing interests declared

### Jon Martin Collinson

- University of Aberdeen, School of Medicine, Medical Sciences and Nutrition, Aberdeen AB25 2ZD, United Kingdom

Contributions: Conceptualisation. Experiment setup and supervision. Experimental data analysis supervision. Wrote the manuscript.

Competing interests: No competing interests declared

### Silke Henkes

- Lorentz Institute, LION, Leiden University, 2311 CA Leiden, The Netherlands

Contributions: Conceptualisation. Model development. Performed numerical simulations. Experimental and simulation data analysis. Wrote the manuscript.

Competing interests: No competing interests declared

## Funding

Jon Martin Collinson - UK BBSRC (grant number BB/J015237/1)

Silke Henkes - UK BBSRC (grant number BB/N009150/1-2), and the support of the NWO sector plan science and Leiden University.

Kaja Kostanjevec - EASTBIO PhD studentship

Rastko Sknepnek - UK EPSRC (Award EP/W023946/1)

## Animal Ethics

Animal experiments were performed under UK Home Office Project Licence number PP5319455 to Jon Martin Collinson, after review and approval by University of Aberdeen Ethical Review Committee.

## Data Availability

A supporting dataset for this publication is available on Zenodo at doi.org/10.5281/zenodo.14261827.

It contains all experimental data from the in-vitro cell cultures and the fixed corneas, as well as the associated analysis software we developed. It also contains the cornea simulation and analysis codes SAMoS and SAMoSA, and example configuration files for cornea and in-vitro type simulations. In addition, we provide sample images and datasets, and the analysis pipeline for the simulations.

## Acknowledgements

We would like to thank Luke Coburn for creating the initial version of the MATLAB stripe fitting algorithm. We thank Luiza Konkina for assistance with tracing corneal striping patterns. RS and SH would like to thank the Isaac Newton Institute for Mathematical Sciences, Cambridge, for support and hospitality during the programme *New statistical physics in living matter*, where part of the work on this paper was undertaken.

---

**Figure 2–Figure supplement 1.**
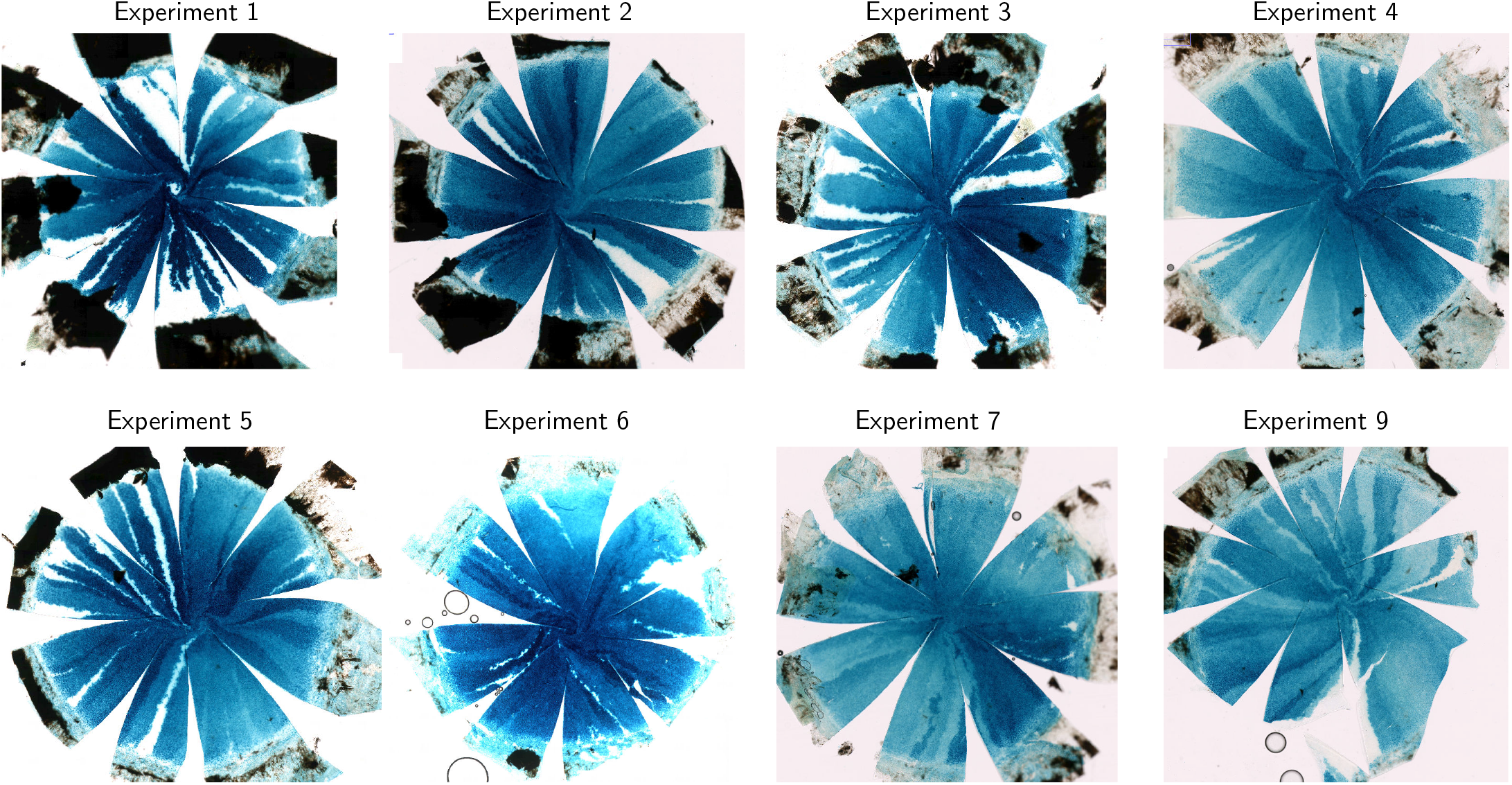
Fixed, stained and dissected LacZ-mosaic corneas (*n* = 8) used for flow inference. The striped corneal surface (white and blue) is flattened by making radial cuts, opening wedge-shaped spaces. The limbal boundary is located at the transition to the brown outer tissue.

**Figure 2–Figure supplement 2.**
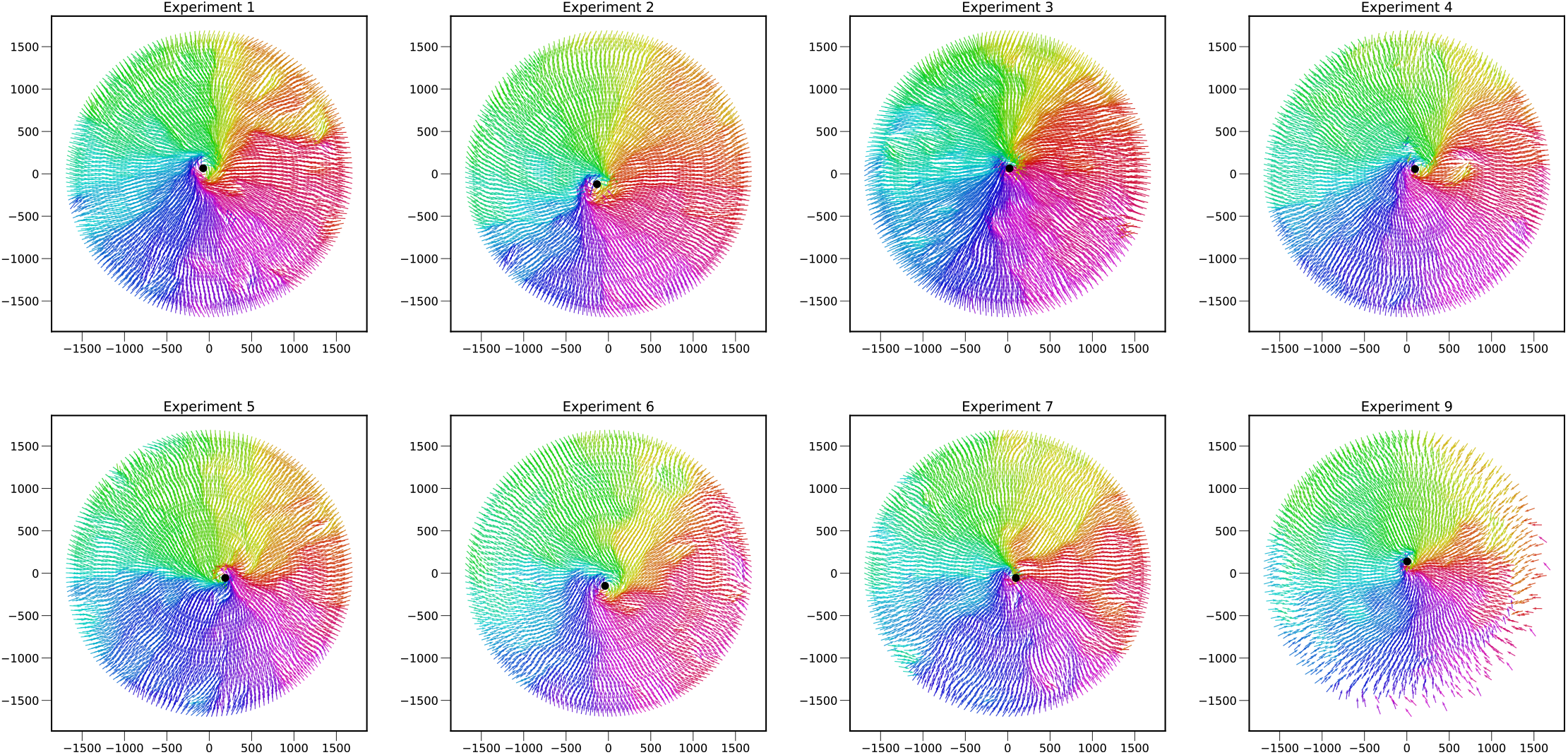
Inferred flow fields for corneas in experiments 1-8.

**Figure 7–Figure supplement 1.**
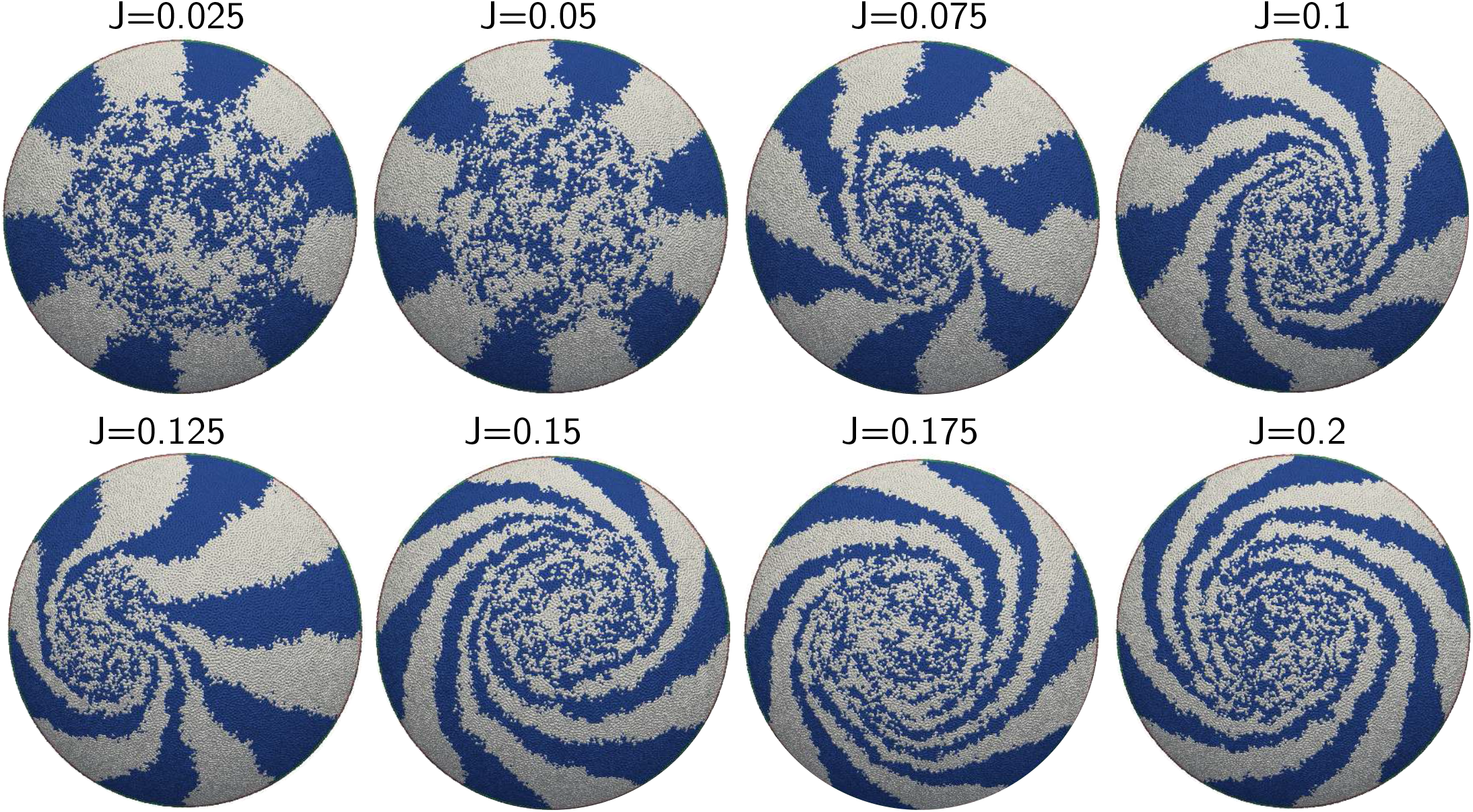
Emergence of the corneal spiral as a function of *J* for *R* = 500 µm simulated corneas, snapshots are at 20 days.

**Figure 7–Figure supplement 2.**
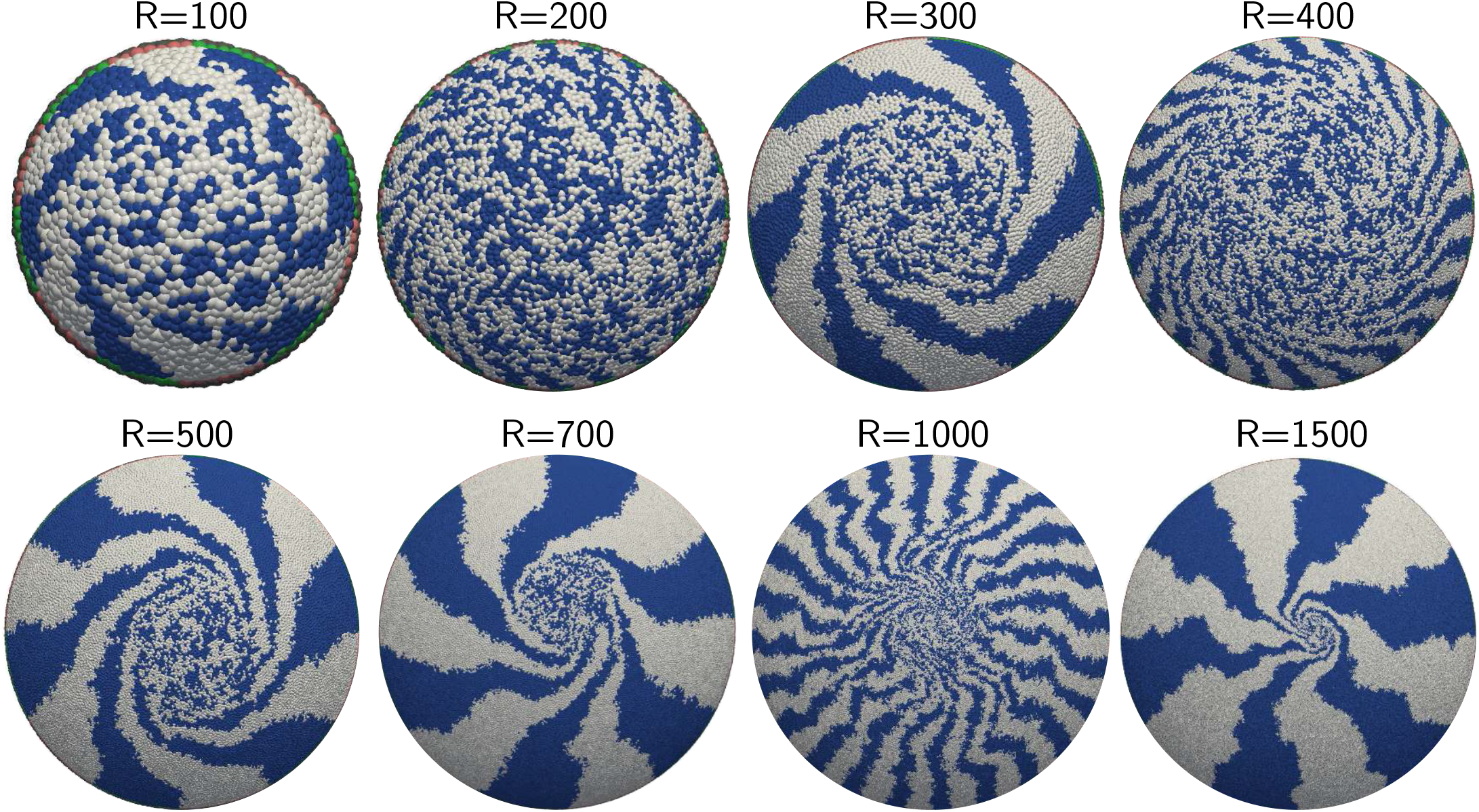
Emergence of the corneal spiral as a function of *R* for *J* = 0.1 h^−1^ simulated corneas, snapshots are at 20 days. Note that simulations for *R* = 200, 400 and 1000 µm have patterned 48 instead of 12 corneal stripes into the limbus.

## Notes

https://zenodo.org/records/14261828

### Competing Interest Statement

The authors have declared no competing interest.

### Summary of Updates

This 2nd revision of the article has been made to comply with the VOR requirements of eLife, where this article will appear shortly.

