## Supplementary material for "A soft active matter models explains spiral epithelial cell migration on in-vivo corneas": Description of supplementary videos

Description of Supplementary videos. Best played with VLC media player.

S1 (S1_explant_wt_5_9.mp4)
Supplements Figure 4 in the paper. Phase contrast image and PIV flow field showing collective cell migration for one corneal explant experiment, labeled 1 in Figure 4.

S2 (S2_plastic_wt_b_24_7.mp4)
Supplements Figure 4 in the paper. Phase contrast image and PIV flow field of corneal epithelial cells grown on tissue culture plastic, showing a flatter phenotype and enhanced migrataion. Experiment is labeled 4 in Figure 4.

S3 (S3_explant_sim_J0.15.mp4)
Supplements Figure 4 in the paper. Simulated matched particle model on the plane for the explant experiments, with arrows representing hydrodynamic velocity, for J=0.15.

S4 (S4_plastic_sim_J0.15.mp4)
Supplements Figure 4 in the paper. Simulated matched particle model on the plane for the tissue culture plastic experiments, with arrows representing hydrodynamic velocity, for J=0.15.

S5 (S5_R100stripes.mp4)
Supplements Figure 5 in the paper. Setup of the corneal model, for R=100 and J=0.1: Blue and white TA cells are derived from green and pink stem cells, respectively.
Stem cells and barrier cells (grey) have fixed positions, and TA cells move over the corneal surface according to eq. 7-8, while dividing and disappearing.
The system quickly reaches a steady state with a fixed density and an emerging spiral flow.

S6 (S6_R700stripes.mp4)
Supplememts Figure 6b in the paper. Emergence of stripes and spiral in a R=700, J=0.1 simulated cornea. The video lasts 20.4 simulated days.

S7 (S7_R700flow.mp4)
Supplements Figure 6c in the paper. Hydrodynamic flow field and flow defects (white = +1, black = -1) for the same R=700, J=0.1 simulation as S2.

S8 (S8_R500_J0.025stripes.mp4)
Supplements Figure 7a in the paper. Absence of spiral formation and of effective corneal resurfacing for J=0.025, below the flocking transition, at R=500.

S9 (S9_R500_J0.025flow.mp4)
Supplements Figure 7a in the paper. Hydrodynamic flow field and flow defects (white = +1, black = -1) for the same R=500, J=0.025 simulation as S4.
The flow field remains disordered with positive and negative defects throughout the simulation.

S10 (S10_R500_J0.2stripes.mp4)
Supplements Figure 7a in the paper. Emergence of stripes and spiral in a R=500, J=0.2 simulated cornea, this spiral is wound more tightly than J=0.1.

S11 (S10_R500_J0.2flow.mp4)
Supplements Figure 7a in the paper. Hydrodynamic flow field and flow defects (white = +1, black = -1) for the same R=500, J=0.2 simulation as S6.
